# Cationic Peptides Facilitate Iron-induced Mutagenesis in Bacteria

**DOI:** 10.1101/026112

**Authors:** Alexandro Rodríguez-Rojas, Olga Makarova, Uta Müller, Jens Rolff

## Abstract

*Pseudomonas aeruginosa* is the causative agent of chronic respiratory infections and is an important pathogen of cystic fibrosis patients. Adaptive mutations play an essential role for antimicrobial resistance and persistence. The factors that contribute to bacterial mutagenesis in this environment are not clear. Recently it has been proposed that cationic antimicrobial peptides such as LL-37 could act as a mutagen in *P. aeruginosa*. Here we provide experimental evidence that mutagenesis is the product of a joint action of LL-37 and free iron. By estimating mutation rate, mutant frequencies and assessing mutational spectra in *P. aeruginosa* treated either with LL-37, iron or a combination of both we demonstrate that mutation rate and mutant frequency were increased only when free iron and LL-37 were present simultaneously. The addition of an iron chelator completely abolished this mutagenic effect, suggesting that LL-37 enables iron to enter the cells resulting in DNA damage by Fenton reactions. This was also supported by the observation that the mutational spectrum of the bacteria under LL-37-iron regime showed one of the characteristic Fenton reaction fingerprints: C to T transitions. Free iron concentration in nature and within hosts is kept at a very low level, but the situation in infected lungs of cystic fibrosis patients is different. Intermittent bleeding and damage to the epithelial cells in lungs may contribute to the release of free iron that in turn leads to generation of reactive oxygen species and deterioration of the respiratory tract, making it more susceptible to the infection.

**Author Summary:** Cationic antimicrobial peptides (cAMPs) are small proteins naturally produced by the immune system to limit bacterial growth mainly through pore formation in the membrane. It has recently been suggested that sub-inhibitory concentrations of cAMPs promote bacterial mutagenesis, similarly to antibiotics. However, we previously reported that cAMPs do not increase mutation rate and do not activate bacterial stress responses. Here we resolve this contradiction. We report that free iron in the culture medium increases mutagenesis in the presence of cAMPs. We show that sub-inhibitory concentrations of cAMPs facilitate entry of free iron into bacterial cells, where it interacts with hydrogen peroxide, thereby resulting in production of DNA-damaging reactive oxygen species and increased mutagenesis. Moreover, these results may have clinically-relevant implications: while very little free iron is normally present in healthy individuals, this is not the case in patients suffering from cystic fibrosis, where elevated bacterial mutagenesis promotes antibiotic resistance and contributes to persistence and severity of infection. Thus, an intervention aimed at reduction of free iron in the lungs could reduce the cAMPs-facilitation of iron-mediated mutagenesis; hence antibiotic resistance and pathoadaptation.

## Introduction

*Pseudomonas aeruginosa* is an important opportunistic pathogen involved in chronic respiratory and hospital-acquired infections [1]. In cystic fibrosis (CF) patients, one of the most common genetic diseases in humans, this bacterium causes chronic lung infections that result in significant morbidity and mortality [2]. *P. aeruginosa* infections are difficult to treat due the inherent resistance to many drug classes, its ability to acquire resistance, via mutations, to all relevant treatments and its high and increasing rates of resistance locally [3].

Mutagenesis plays a crucial role in adaptation of this pathogen for persistence and antibiotic resistance acquisition in CF [4], being found a high proportion of hypermutable bacteria among *P. aeruginosa* isolates [5]. Recently, Limoli *et al* [6] reported an increased mutant frequency after treatment of *P. aeruginosa* with the human cationic peptide LL-37. Based on this finding they proposed that cationic peptides elevated bacterial mutation rates. In a recent study, we reported mutation rates of *E. coli* in the presence of cationic antimicrobial peptides (cAMPs) and antibiotics. LL-37 was present in the panel of AMPs that we tested and we did not find any increase in mutation rates. We also used transcription reporter assays and qRT-PCR and showed that none of the AMPs elicited the main mutagenic stress pathways of bacteria SOS or *rpoS* [7].

Here, we aim to resolve this apparent contradiction based on the observation that the different studies used different media. Limoli *et al*. [6] used M63 for *P. aeruginosa* and LB for *E. coli*, while we used non-cation adjusted Mueller-Hinton Broth (MHB), commonly used for cAMP susceptibility testing. The most striking difference in culture media between the two studies is that both, M63 [8] and LB are iron-rich [9,10], while MHB is not [11]. Fe^2+^ catalyses hydroxyl radical formation by reacting with hydrogen peroxide both intra- and extra-cellularly, the Fenton reaction [12]. Without free iron hydrogen peroxide reactivity is low at physiological pH and iron metabolism is strictly controlled to avoid DNA and other damage caused by oxygen radicals [13,14]. In most natural situations iron is in short supply, but in cystic fibrosis [15,16] due to accult bleeding of highly vascularised lung tissue and haemoptysis particularly during acute exacerbations, and damage of the respiratory tract epithelium where Fe^2+^ is present intermittently. Although ferrous iron is prone to oxidation, the anaerobic growth of at to high density of bacteria [17] may contribute to stabilise this metal in the reduced state.

Many cationic antimicrobial peptides change membrane permeability properties [18], and this led us to hypothesize that sub-inhibitory concentrations of LL-37 increase uncontrolled iron transport from the extracellular space to the cytoplasm without the intervention of the bacterial iron trafficking system. An intracellular surplus of iron should then result in DNA damage caused by the Fenton reaction.

Here, we (i) estimate the mutation rate of *P. aeruginosa* in the presence of LL-37, iron or both; (ii) we then investigate the hypothesis that ferrous iron (Fe^+2^) is causal to an increase in mutation rate and that LL-37 facilitates this process, something that is true for colistin too; and finally (iii) we investigate the mutational spectra to find out if promoted mutations match with any of the molecular signatures of Fenton reactions.

## Results and Discussion

### Mutation rate is increased in the presence of ferrous Fe^2+^ and LL-37

First we determined the mutation rate of *P. aeruginosa* using a fluctuation assay, in the presence of either LL-37 (32 μg/ml), Fe^2+^ (40μM), or both (figure 1). Mutation rate was only increased when LL-37 and Fe^2+^ were added simultaneously, while addition of LL-37 or Fe^2+^ separately did not produce the same effect. To investigate the effect of different concentrations of LL-37 and Fe^2+^, we estimated mutant frequencies. None of the LL-37 concentrations tested, ranging from 4 to 32 μg/ml, yielded any detectable changes in mutant frequency (figure S1). Fe^2+^ alone, at different concentrations, also did not increase the mutant frequencies in comparison with the control group (figure S2).

**Figure 1.**
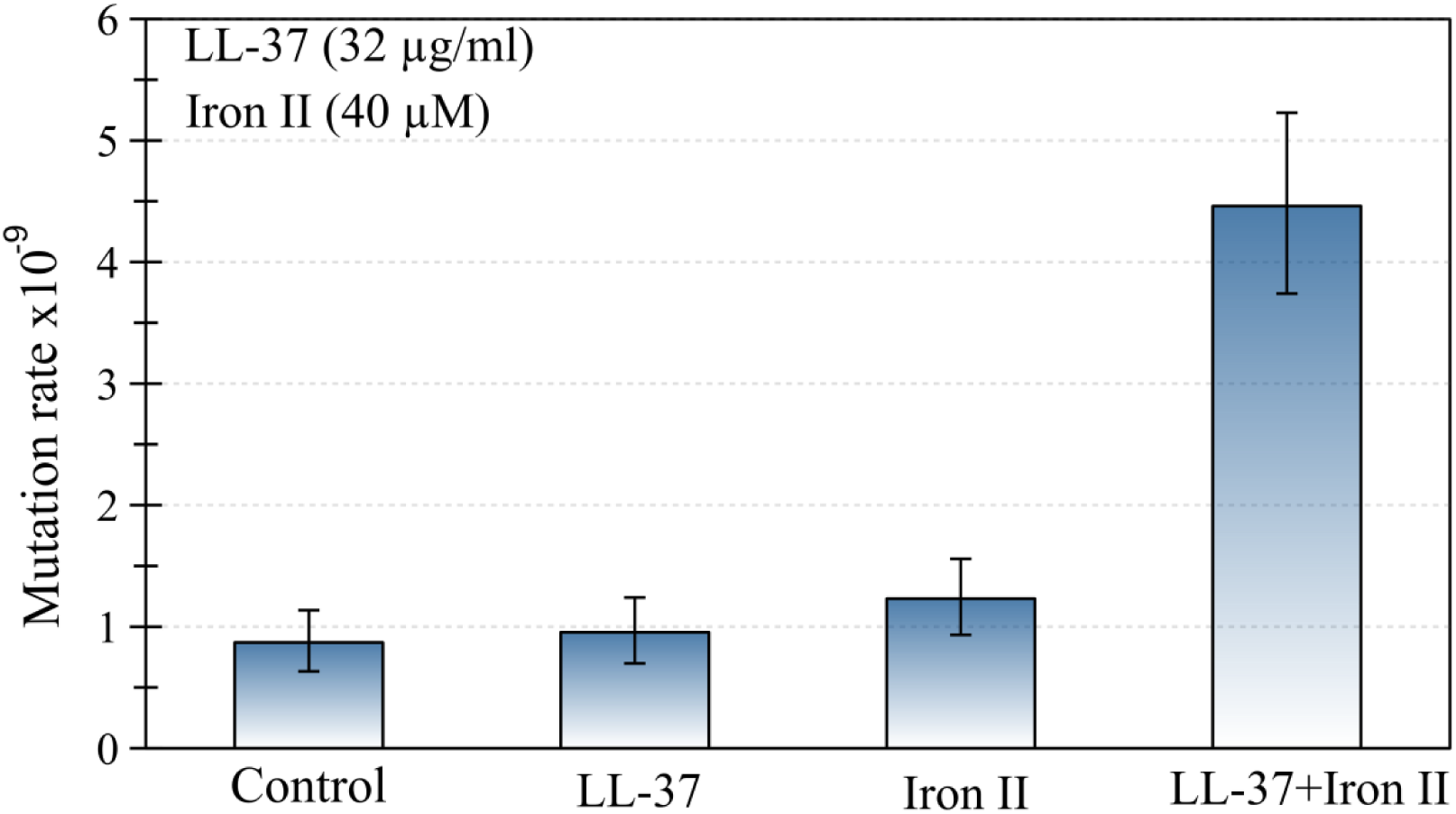
The joint action of LL-37 and ferrous ions induces an increase in the mutation rate of *P. aeruginosa.* No changes occur when LL-37 or iron are added to the culture separately. Error bars represent 95 % confidence intervals for mutation rates.

Subsequently, we assayed the MIC50 (32 μg/ml) of LL-37 with several iron concentrations and measured the impact on rifampicin mutant frequency of *P. aeruginosa* strain PAO1. We found that added concentrations of 10, 20, 40 and 80 μM of Fe^2+^, increased the mutant frequency between three to five times (figure S3). Limoli *et al.* [6] observed increased mutant frequencies in both, *P. aeruginosa* and *E. coli*, proposing that LL-37 induces mutagenesis in these bacteria. Taken at face value, this contrasts with our work, where we did not find such effect in *E. coli* [7]. *P. aeruginosa* in the Limoli *et al.* study however, was cultured in an iron-rich medium, M63 [8], which contains 0.5 mg of iron sulphate per litre (Fe^2+^). To confirm the results in a different bacterial model, the experiment was repeated with *E. coli* using LB as a culture medium. However, LB is also iron-rich (∼16 μM of iron) [9,10]. In these experiments, just before the treatment with LL-37 in a saline solution, bacteria were washed. It has, however, been shown that Gram-negative bacteria can actively accumulate iron in the periplasmic space [19]. This led us to hypothesise that sub-lethal concentrations of cationic antimicrobial peptides can facilitate iron transport into the cells.

### LL-37 facilitates mutagenesis but iron is causal

To confirm that iron is causing the observed increase in mutagenesis, we first repeated the assay in the presence of an iron chelator, 2-2′ bipyridyl. 2-2′ bipyridyl completely abolished the increase in mutant frequency in the tested concentration of LL-37+Fe^2+^ combination. The sole action of the chelator plus Fe^2+^ did not increase the mutant frequency (figure 2).

**Figure 2.**
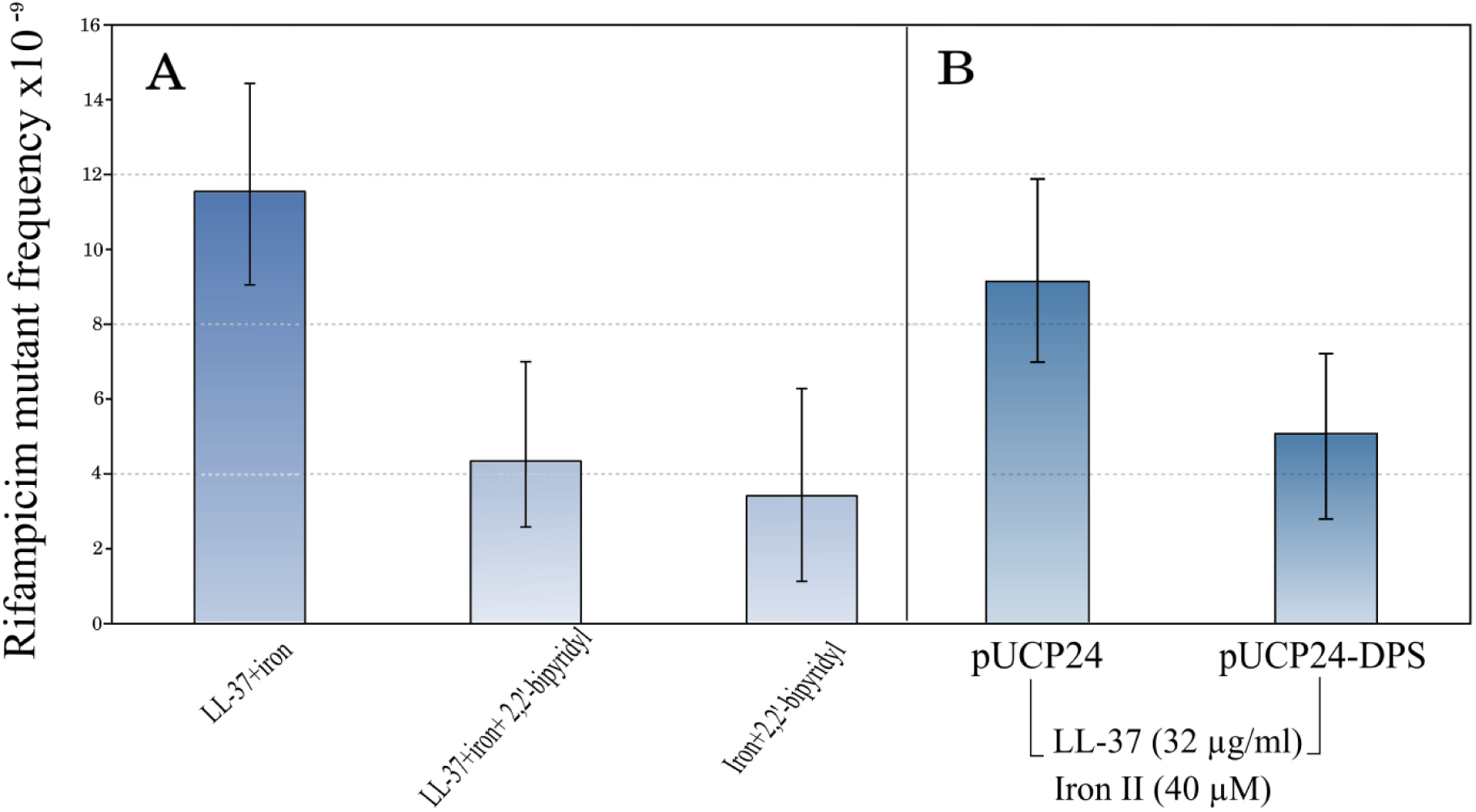
The mutagenesis of LL-37+Fe^2+^ combination is suppressed by the addition of an iron chelator. This supports the notion that iron is causal in increasing mutagenesis (A). In the same line, over-expression of Dps, a natural iron chelator in bacteria, also decreases the mutant frequency to rifampicin (B). Error bars represent 95 % confidence intervals for mutant frequencies.

Bacteria have a number of mechanisms to keep free iron as low as possible inside the cell. One of these is Dps, a natural iron chelating protein. To confirm the results obtained with 2-2′ bipyridyl, we used a Dps over-expressing strain by cloning the *dps* gene into a multi-copy plasmid under a constitutive promoter. The mutagenesis was not completely reverted in comparison with non-treated controls, but there was a 1.8 fold-reduction (figure 2).

The iron uptake assay showed that there was a significant difference (*P*=0.00015) in total iron content in bacterial cells between LL-37+ Fe^2+^ (17.50 ± 5.10 nmoles/ml, mean ± standard deviation) and iron alone (5.65 ± 4.13 nmoles/ml, mean ± standard deviation) treatments after 30 minutes of incubation, indicating that LL-37 indeed promotes a non-physiological free iron entrance.

We also tested a further AMP: Colistin, which is of bacterial origin, and is one of the most effective agents against *P. aeruginosa* in CF infections [20]. We treated cultures of *P. aeruginosa* PAO1 with MIC50 of colistin (1.8 μg/ml) in high and low concentrations of iron in the media. The effects of colistin were very similar to the effects we found for LL-37 (figure 3). Although the mechanism of action of colistin is not fully understood [21], our results suggest pore forming mechanism or permeability changes in the cell as likely mechanisms. It has been demonstrated that colistin promotes the uptake of hydrophilic antibiotics, explaining their synergism with them [22].

**Figure 3.**
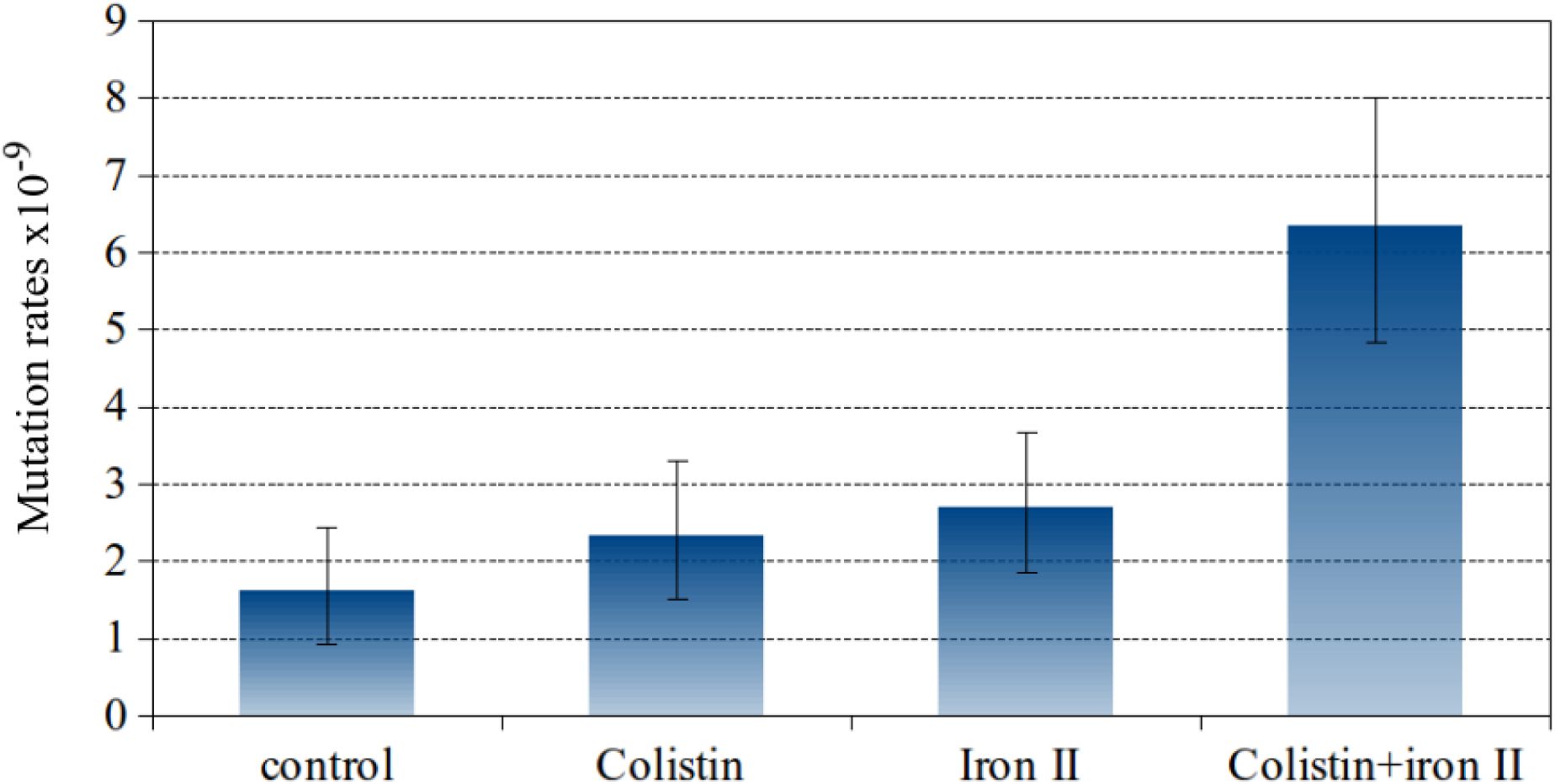
Mutagenesis induced by colistin and Fe^2+^ combination. The mutation rate is only increased if both, iron and colistin, are present. Error bars represent 95 % confidence intervals for mutation rates.

Previous work has suggested that iron is involved in *Pseudomonas aeruginosa* pathoadaptation and antibiotic resistance acquisition [15]. All experiments were carried out with the more soluble Fe^2+^ because ferrous iron is abundant in the CF lung (∼39 μM on average for severely ill patients) and significantly correlates with disease severity [23]. Moreover, iron activates the two-component signal transduction system BqsRS in *P. aeruginosa,* which is transcriptionally active in CF sputum, and promotes tolerance to cationic stressors [24]. This increases the tolerance to both host peptides such as LL-37 and colistine, which is administered in CF therapy.

The new mechanism proposed here has almost certainly important implications. Under specific pathologies such as cystic fibrosis or other respiratory chronic diseases, iron that is usually in short supply, is abundant. Disturbances in iron metabolism have been shown to promote evolution of antibiotic resistance in *E. coli* [25]. Given that cationic antimicrobial peptides usually induce membrane permeability in all susceptible microbes and Fenton chemistry is universal, the mechanism we propose here is likely applicable to other AMPs. For example, beta-defensin-2 was shown to double the mucoid conversion rate, a mutagenic process, of *P. aeruginosa* [6]. Yet free iron is rarely available in most physiological situations but for a few pathologies such as cystic fibrosis

Additionally, it has been reported that some cationic antimicrobial peptides interact with bacterial membrane proteins and delocalise them [26]. It is conceivable then that LL-37 may interact and interfere with iron transport systems, which in turn may contribute to iron homeostasis disruption and enhance mutagenesis. This possibility requires additional investigation and will be the goal for future studies.

In the context of cystic fibrosis, the increase in salt concentration may lead to the reduced activity or complete inactivation of other antimicrobial peptides, as observed for human beta-defensin-1 [27], while other components could eventually contribute to mutagenesis in the same way that LL-37 does. Moreover, the PhoP-PhoQ and PmrA-PmrB two-component regulatory systems of *P. aeruginosa* may play an important role in antimicrobial peptide tolerance. This resistance is reproduced *in vitro* when magnesium concentrations are low [28]. Our experimental conditions, where divalent cations are depleted or in low concentrations, seem comparable. In the light of our results this suggest that bacteria under certain conditions that elicit the expression of these two component systems alter the lipidA structure resulting in increased resistance to colistin [29]. The same systems up-regulate ferrous iron uptake which is mediated by feoAB operon [28,30]. This phenomenon could potentially contribute to saturate intracellular iron storage systems and to generate an excess of iron that eventually can participate in Fenton reaction operating by the mechanisms that we propose here.

How would the mechanism we propose here enhance the overall mutation rate of bacterial populations? In the context of cystic fibrosis, there is a high proportion of hypermutable bacteria due to the inactivation of their DNA miss-match repair (MMR) genes [31]. We may expect a synergistic effect of both types of mutagenesis as we proposed in the past for the mutagenic effect of cystic fibrosis lung environment and the intrinsic mutagenesis of *P. aeruginosa* [4]. It can be speculated that iron-mutagenesis can facilitate the rise of mutator bacteria by enhancing the inactivation of MMR genes. This could then weaken genetic constraints that impede the evolution of bacteria to resist antibiotics by multiple pathways as previously described [32].

### Mutational spectra of iron-induced mutagenesis

The results above strongly suggest that sub-lethal concentrations of cationic antimicrobial peptides facilitate the access of free iron to the cytoplasmic compartment of *P. aeruginosa.* Given that the Fenton reaction results in DNA damage [33] and reactive oxygen species (ROS) damages to the DNA display specific molecular fingerprints, we investigated if there was evidence of the Fenton reaction effects on the mutational spectrum.

It would be very difficult to investigate the mutational spectrum in *rpoB* (sub-unit of RNA polymerase) that confers resistance to rifampicin in *P. aeruginosa* PAO1. This gene is essential in and detectable mutations are mostly point mutations that would constrain the analysis to a few types of changes. We therefore decided to use another strain, PA14, where mutations that confer resistance to fosfomycin are well characterized [34–36]. Resistance mutations to fosfomycin in *P. aeruginosa* PA14 depend on a single non-essential locus encoding the glycerol-3-phosphate transporter GlpT, making it a much better marker for studying the entire repertoire of mutations compared to *rpoB* [36,37] [38].

The treatment of *P. aeruginosa* P14, which shows a similar sensitivity to LL-37 as the PAO1 strain used above, with the mutagenic combination of LL-37+Fe^2+^ showed almost a four fold-increase in mutant frequency (figure S4).

To assess how addition of free iron and antimicrobial peptides affect the mutation spectrum of the *glpT* gene of PA14 strain we exposed bacteria to either free iron, antimicrobial peptide LL-37, or both. All of the resulting resistant mutants contained non-synonymous substitutions or deletions in *glpT* that potentially affect the stability of GlpT transporter and likely disrupt the function of the protein (figure 4, table S1, figures S5 and S6). LL-37+Fe^2+^-treated bacteria displayed striking differences in mutational spectra when compared to the other treatments (figure 4 and table S1). We used Monte Carlo hypergeometric test implemented in iMARS, a mutation analysis reporting software [39], to assess the overall differences between mutational spectra. The probability of the mutational spectra to be the same stood at 0.554706 (*P*-value confidence limits 0.531080 – 0.578) when the iron treatment was compared to LL-37, but was below 0.0000001 when either were compared with LL-37+Fe^2+^ treatment. Moreover, we found a mutation hotspot (R93 to W change, 10/20 clones) in the LL-37+Fe^2+^ treatment group that was significantly (*P*=0.0004, two-tailed Fisher’s exact-test) different from the two other treatments (tables S2, S3 and S4). This mutation hotspot is a single nucleotide transition from C to T, which is one of the most frequent types of mutations caused by ROS [40]. We found that twelve out of twenty clones from the LL-37+Fe^2+^ condition had C to T transitions, whereas none were present in either iron or LL-37-treated bacteria (two-tailed Fisher’s exact-test (*P*< 0.0001) (figure S7). It is striking that although several C to T mutations can potentially lead to *glpT* inactivation are in principle possible, that the majority are concentrated in a single point. Such mutation hotspots can be driven by a specific topology on the chromosome [41], or a particular sequence prone to double strand breaks resulting in mutations after repair [42]. In general, observable mutations are the results of mutation-repair balance and not all mutations are repaired with the same efficiency. A good example in *P. aeruginosa* is the mutational inactivation of the anti-sigma factor gene, *mucA*, with the mutated allele *mucA*22 most prevalent (25 to 40%). This inactivation seems to be spectrum dependent [43].

**Figure 4.**
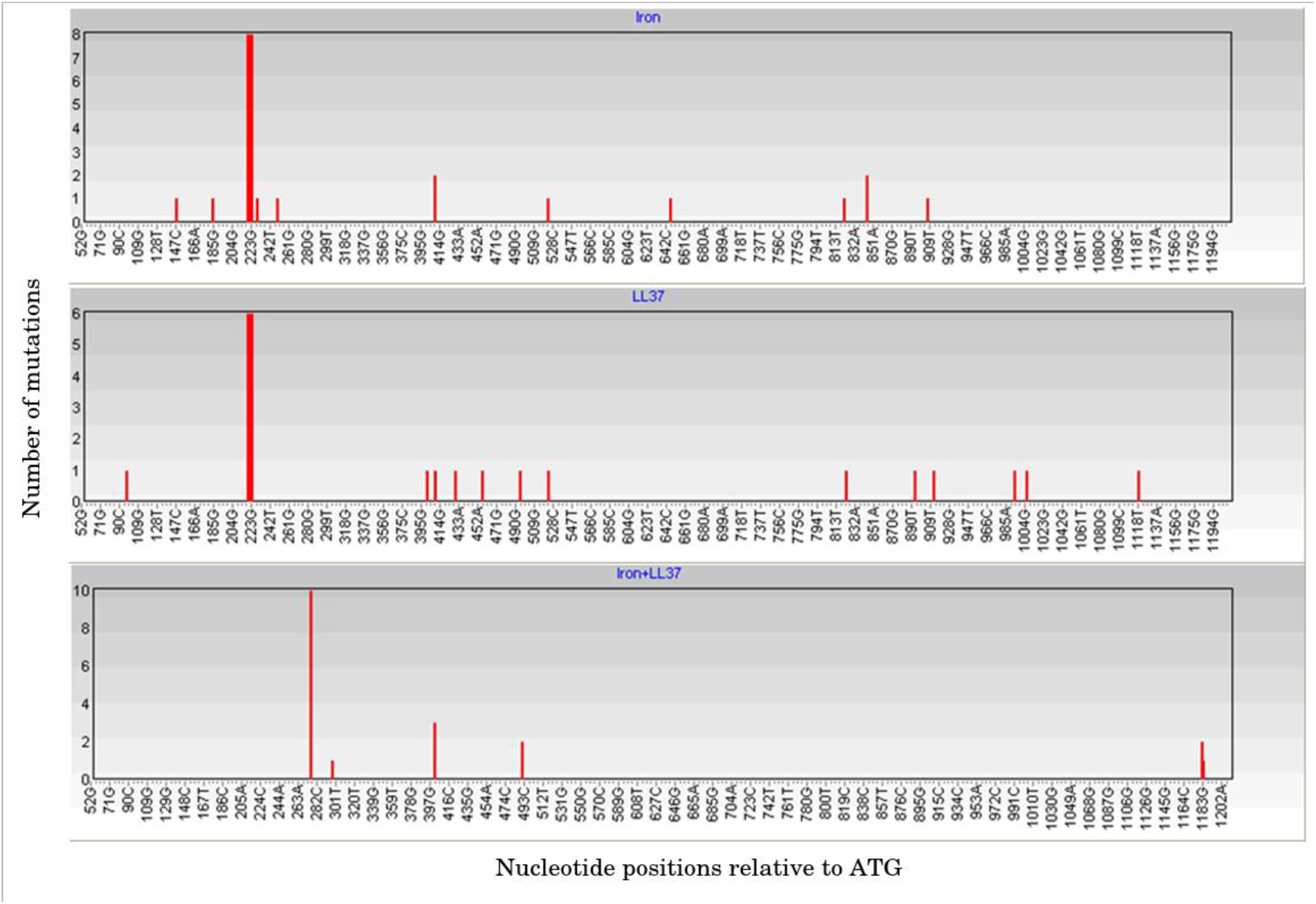
Mutation hotspots in *glpT*. Distribution of the mutations in the 1160 bp-long fragment of *glpT* in *P. aeruginosa* PA14 fosfomycin-resistant clones treated with iron, LL-37, or a combination of both. The number of mutations (nucleotide substitutions, indels) is plotted against respective nucleotide positions within the gene fragment. Note the overlapping mutations at positions 220-225, 409 and 524 bp in iron and LL-37 treatments and absence of common mutations in LL-37+Fe^2+^ treatment. The figure was generated using the mutational spectrum analysis software iMARS [39].

Interestingly, C to T transitions were one of the most frequent types of single nucleotides changes in the genomes of *Salmonella typhimurium* evolved to increasing concentrations of LL-37 in modified LB medium short of sodium chloride and anions [44], which is fully consistent with our results.

A number of studies [45–48], which caused some controversy [49–51], suggested that hydroxyl radicals can be generated as a consequence of antibiotic treatments and this aggressive by-product may take part in the killing mechanism of bactericidal drugs or promote mutagenesis. Most of these studies were carried out in the iron-rich LB medium and whether processes as described here for the interaction between an AMP and iron apply to antibiotics remains to be explored.

### Conclusions

Our results support the notion that under certain pathological situations, sub-inhibitory concentrations of cAMPs facilitate uncontrolled uptake of free iron by bacterial cells, which results in increased mutagenesis by Fenton reaction (figure 5). According to our results, this could be a general mechanism underlying mutagenesis by joint action of antimicrobial peptides and free iron in specific situations where iron is not limited.

**Figure 5.**
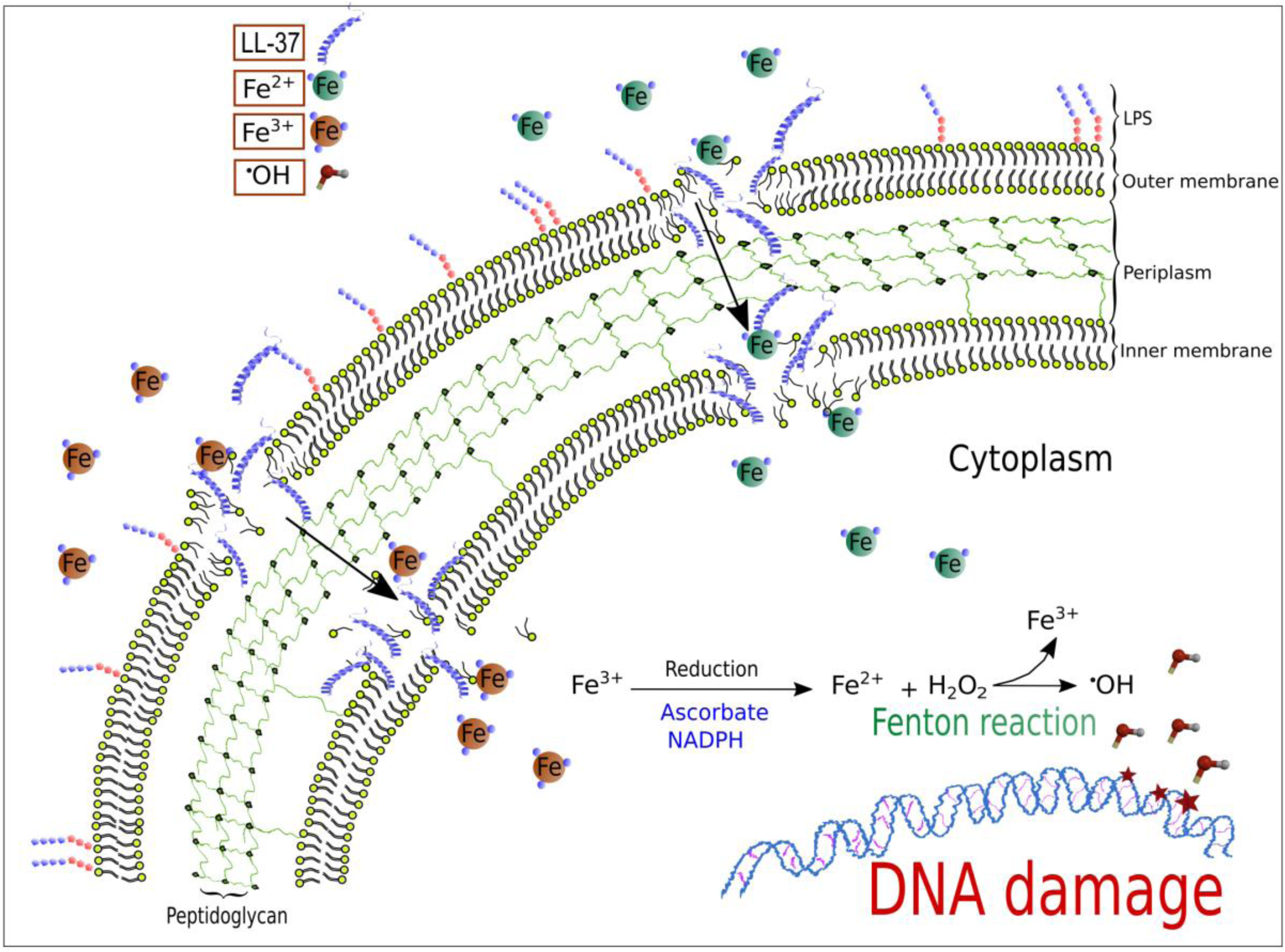
A model for LL-37-mediated, iron–induced mutagenesis in *Pseudomonas aeruginosa*. The interactions of LL-37 with the membrane at sub-inhibitory concentrations lead to transient permeability changes in the membranes that promote iron movement in favour of the electrochemical gradient. Uncontrolled uptake of ferrous ions stimulates Fenton reaction that leads to hydroxyl radical formation and results in DNA damage and mutagenesis.

Free iron levels are kept as low as possible due to its toxic effect for all living beings but especially in bacteria that lack proper cell compartments. In fact, iron withdrawal is part of the natural innate immune response in inflammation that makes free iron even more scarce. During inflammation and infection a “hypoferremic response” (anaemia of inflammation) is observed [52,53]. Many chelating proteins such as transferrin and ferritin exhibit antibacterial activity simply by making iron availability incompatible with bacterial proliferation [54].

Despite showing that many cationic antimicrobial peptides are unable to increase mutation rates [7], the particular situation of iron-induced mutagenesis can be of great interest for certain types of infections where iron homoeostasis is compromised. In cystic fibrosis, bleeding happens frequently. Considering that *P. aeruginosa* is one of the most common pathogens that acquire all antibiotic resistance by mutations, the mechanism proposed here is likely very relevant for pathoadaptation.

Finally, our work has potential implications for the development of future treatment of chronic respiratory infections by *Pseudomonas aeruginosa.* For example, modulation of iron-chelating agent in CF therapy could potentially slow down the pathoadaptation and development of resistance in CF, and diminish lung damaging by ROS.

## Materials and methods

### Bacteria and growth conditions

The *P. aeruginosa* PAO1 wild-type strain was kindly provided by M. A. Jacobs. The *P. aeruginosa* PA14 was kindly provided by Nicole T. Liberati and Frederick Ausubel. All bacterial strains were cultured in Mueller-Hinton Broth, non-cation adjusted (Sigma), with total iron content 0.22 ± 0.02 μM, following recommendations of CLSI for cationic antimicrobial peptide susceptibility testing. The MHB pH was adjusted to 6 in all cases with acetic acid to enhance solubility of both, LL-37 and iron compound. All experiments were performed at 37°C, under agitation in liquid culture. For genetic manipulation, *Escherichia coli* DH5α strain was used and routinely cultured in Lysogeny Broth (LB medium), supplemented with antibiotic when appropriate.

### Minimal inhibitory concentration (MIC)

MICs were determined according to CLSI recommendations by a microdilution method with some modifications for antimicrobial peptides. The MIC was defined as the antimicrobial concentration that inhibited growth after 24 hours of incubation in liquid MHB medium at 37°C. Polypropylene non-binding multi-well plates (Th. Geyer, Germany) were used for all experiments.

### Determination of MIC50

The MIC50s for all antimicrobials were determined by inoculating strains grown to mid-log phase into the wells of a 96-microwell plate. Approximately 10^2^ cells from overnight cultures of PAO1 and PA14 strains were inoculated into 50 ml tubes containing 10 ml of MBH and incubated at 37°C with strong agitation until the mid-log phase of growth (approximately 108 cfu/ml). Then, 100 μl of 2 ×10^8^ cells from these cultures were inoculated in each well of polypropylene non-binding 96-multiwell plates containing 100 μl of fresh Mueller-Hinton medium with growing concentration of serially diluted LL-37. The plates were incubated at 37°C during four hours with continuous agitation in a plate reader (Synergy HT, BioTeK). Four replicates per concentration were prepared and the experiments were repeated twice. MIC50s at 4 hours were defined as the concentrations at which 50% of growth reduction in comparison to the control at OD_600_ were observed.

### Estimation of mutant frequencies and mutation rates

For spontaneous-mutation rate measurements of PAO1 strain, 1/100 dilutions of overnight cultures were inoculated into four tubes per group, each containing two ml of MHB medium. The cultures were incubated at 37°C with strong agitation to reach ∼10^8^ cfu/ml. At this point, appropriate concentration of LL-37, colistin, iron sulphate (FeSO_4_) or combinations of antimicrobial peptides and iron, were added to the cultures. The tubes were allowed to continue their normal growth overnight until saturation. In the experiments with colistin, the bacterial suspensions were washed twice with saline solution 0.9 % NaCl before plating. The cultures were appropriately diluted and plated on MHB agar plates with or without rifampicin (100 μg/ml). The mutant frequency was estimated by the number of colonies growing on rifampicin divided by the number of total cfu/ml. To confirm the results, relevant concentrations were also assayed with ten replicates to see the influence of the treatment on the population mutation rates (the number of mutations per cell per generation). Mutation rates were calculated by maximum verisimilitude method and data were processed using the on-line web-tool Falcor (http://www.mitochondria.org/protocols/FALCOR.html) as recommended [55,56]. Falcor software was used to estimate the mutant frequency too. The mutation rates for the strain PA14 under the selected treatments were determined in the same way as described for PAO1, but fosfomycin (128 μg/ml) was used instead of rifampicin.

### Sequencing of fosfomycin resistant mutants (Fos-R) and sequence analysis

To assess the effects of iron and antimicrobial peptides and their combination on the mutation spectrum of *P. aeruginosa* strain PA14, the *glpT* gene of twenty randomly selected Fos-R clones from independent cultures for each treatment group (MHB supplemented either with 40 μM Fe^2+^, LL-37 (32 μg/ml) or a combination of LL-37+Fe^2+^ at same concentrations for both), was amplified by colony PCR using *glpT*-P14-F1 (5-AGCGGAGCTCGCGATGTTC-3) and g*lpT*-P14-R1 (5-TCAGCCGGCTTGCTGCGG-3) primers [36] and Kapa2G Fast ReadyMix PCR with dye kit (KAPA Biosystems, Boston, US). Cycling conditions were as follows: 95°C 7’/(95°C 15’’/60°C 15’’/72°C 40’’)x 35/72°C 7’/4°C hold. The PCR products were purified and sequenced at Macrogen Europe using the forward and reverse primers described above. Sequences were assembled using SeqTrace software. Assembled sequences were imported into CLC Sequence Viewer 6 and aligned using default settings. Low quality flanking sequences were removed and the alignment was trimmed to the 1160 bp fragment (the 52-1212 bp region relative to the A in the start ATG codon of the 1347bp-long *glpT* ORF). A tridimensional homology model of GlpT was generated using Cn3D software by performing a BLASTP search using PA14 strain GlpT protein sequence as a query and mapping the resulting alignment against the experimentally determined *Escherichia coli* K-12 GlpT protein structure. To evaluate the potential effects of amino acid substitution on protein stability, the online tool I-Mutant (http://gpcr2.biocomp.unibo.it/cgi/predictors/I-Mutant3.0/I-Mutant3.0.cgi) was used. TMHMM server v. 2.0 (http://www.cbs.dtu.dk/services/TMHMM/) was used for prediction of transmembrane helices in *glpT* protein sequence. Mutational spectrum differences were analysed using the software iMARS [39].

### Influence of 2-2′ bipyridyl on LL-37+Fe^2+^ mutagenesis

The effect of 2-2′ bipyridyl, an iron chelating agent, on LL-37+Fe^2+^ mutagenesis was determined by measuring its influence on mutant frequency on a selected concentration of LL-37+Fe^2+^ combination (32 μg/ml and 40 μM of Fe^2+^respectively), where mutagenesis was observed. The experiment consisted of adding a titrating concentration of 2-2′ bipyridyl (114 μM) to chelate 95 % of the added iron of treated cultures, in order to make the treatment compatible with bacterial growth. Cultures with the described LL-37+Fe^2+^ combination with no addition of 2-2′ bipyridyl were used as a control. The mutant frequencies of both groups were determined as described elsewhere in this section. LL-37, iron and 2-2′ bipyridyl were simultaneously added to the exponentially growing cultures.

### Cloning of *dps* gene and mutagenesis experiment

DNA fragment containing the PAO1 *dps* gen from genomic DNAs was amplified by PCR using the oligonucleotides PA-DPS-F1 (5′-ATGGAAATCAATATCGGAATCG-3′) and PA-DPS-R1 (5′-CTACTCAAATCAAGCGGTTGGC-3′) as forward and reverse primers, respectively. The fragment contains the ATG codon and 50 nucleotides downstream of the stop codon. The PCR product was directly cloned into the *Sma*I-digested and T-tailed pUCP24 plasmid vector (replicative in both *P. aeruginosa* and *E. coli*), which harbours Gentamicin resistance markers [57]. *E. coli* DH5α was used following standard protocols for genetic manipulations. The resulting plasmids, termed pUCP24-DPS, were introduced by electroporation into PAO1 wild type strain. The cloning vector was also transformed into the same strain as control. An experiment similar to the one designed for 2-2′ bipyridyl was carried out. A mutagenic combination of LL-37+Fe^2+^ was assayed in the strains carrying pUCP24-DPS plasmid or the empty vector pUCP24 and mutant frequencies were determined for both groups.

### Quantification of iron concentrations

Total iron quantification was carried our as previously described with minor modifications [58,59]. Cultures of *P. aeruginosa* PAO1 were grown to an OD_600_ of approximately 0.5 at 37°C with agitation in a volume of two ml in MHB. The cultures were centrifuged at 4000 *g* during ten minutes at 20°C. The pellets were re-suspended in fresh MHB and three different groups were prepared. The treatments consisted of LL-37 (32 μg/ml) (I), iron sulphate to a final concentration of 40 μM (II), a combination iron sulphate and LL-37 (III), both of them at the same concentrations of their respective group I and II, and a control group (IV) to which the proportional amounts of LL-37 and iron sulphate solvent were added (sterile dH_2_0 and dH_2_0, pH=5 respectively). The cultures were incubated for up to 30 minutes and harvested by centrifugation as before but at 4°C. The cell pellets were washed twice with ice-cold phosphate-buffered saline (PBS) and re-suspended in 1 ml of TE buffer containing 5 mg/ml of egg lysozyme (Sigma) and incubated during 10 minutes at room temperature. To quantify total iron, the lysate (one ml) was mixed with one ml of HCl 10 mM and 1 ml of iron-releasing reagent containing HCl 1.4 M + 4.5% (weight/volume) aqueous solution of KMnO_4_; 1/1 and incubated at 60°C for two hours. After cooling, 0.06 ml iron-detection reagent (6.5 mM ferrozine, 6.5 mM neocuproine, 2.5 M ammonium acetate, 1 M ascorbic acid in water) was added and the sample absorbance was read at 550 nm in a plate reader Synergy HT (Biotek). The iron concentrations were determined based on a standard curve obtained with increasing concentrations of ferric chloride and normalized to protein concentration of the lysates. Each group consisted of five cultures. We determined the content of total iron in our cultures media MHB and LB, using the same procedure describe above, starting by the addition of 1 ml of HCl 10 mM and 1 ml of the iron-releasing reagent. Under the suspicion that MHB had lower iron content, the samples of this medium were prepared ten-fold concentrated.

### Statistical analyses

An unpaired Student’s t test or Mann-Whitney U test was used where appropriate for statistical analysis, according to the nature of the data (parametric or nonparametric adjustment). Two-tailed Fisher’s exact test or Monte Carlo hypergeometric test were used to calculate statistical significance of differences in mutation frequencies at each codon site of the alignment between treatments in the mutational spectrum analysis. P values less than 0.05 were considered statistically significant. All tests were performed with statistic software R except for mutational spectrum analysis where iMARS [39] was used instead.

## Acknowledgments

We are grateful to Paul Johnston for comments on the manuscript.

## Supplement figure and table legends

**Figure S1.**
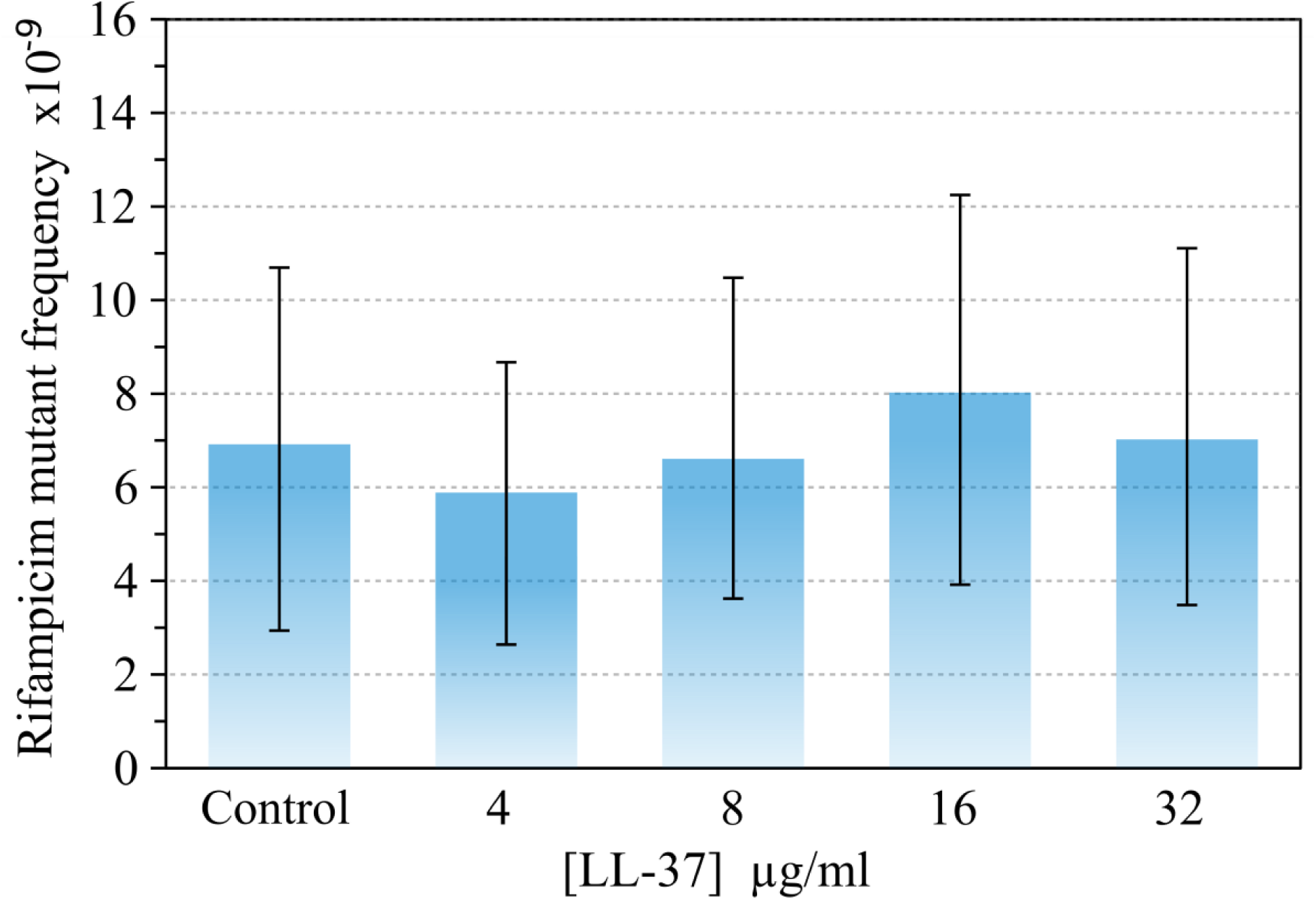
Different concentrations of LL-37 (from 4 to 32 μg/ml) have no impact on mutant frequencies to rifampicin of *P. aeruginosa* PAO1 in MHB. Error bars represent 95 % confidence intervals for mutant frequencies.

**Figure S2.**
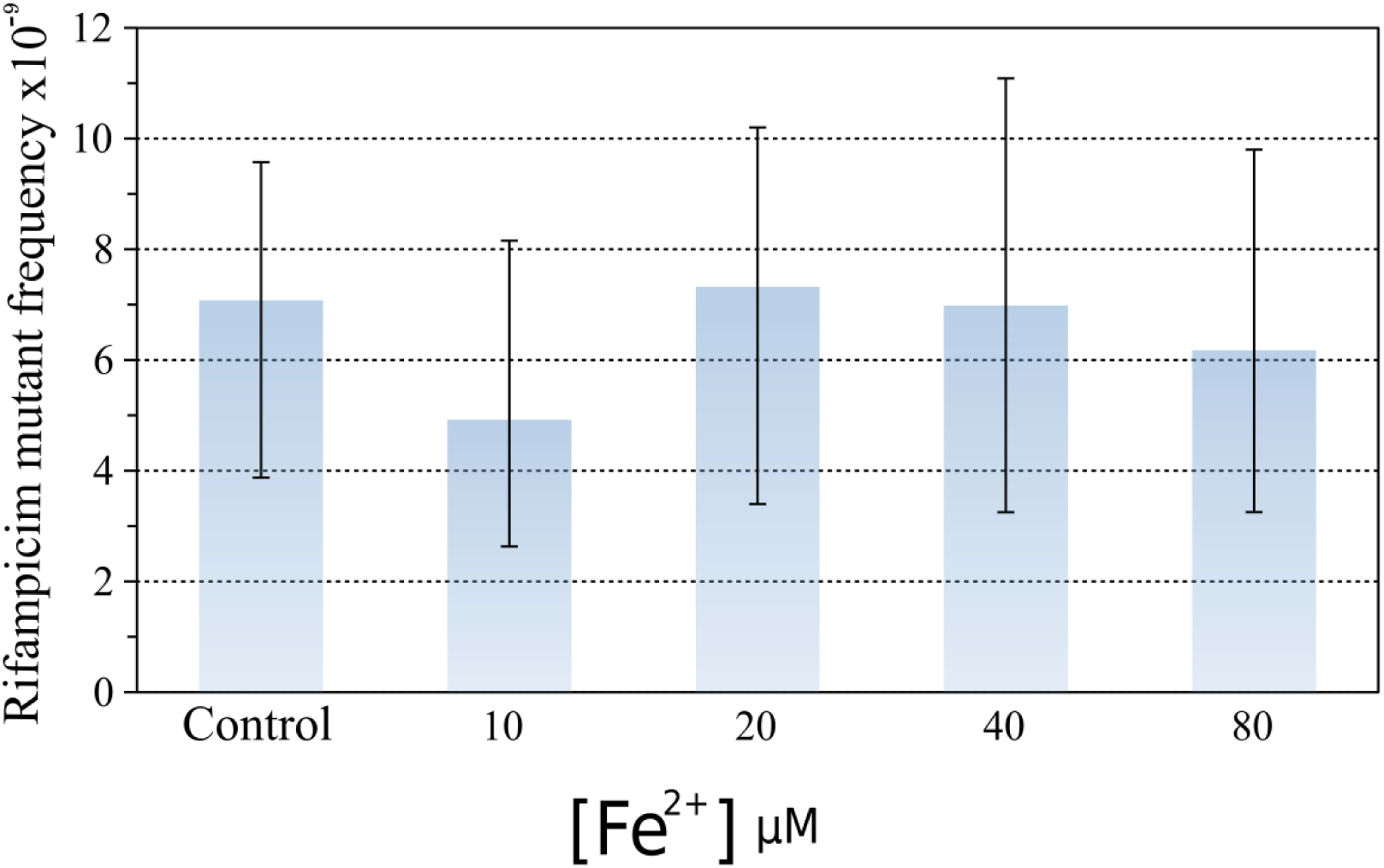
Different concentrations of Fe^2+^ have no impact on mutant frequencies to rifampicin of *P. aeruginosa* PAO1. Error bars represent 95 % confidence intervals for mutant frequencies.

**Figure S3.**
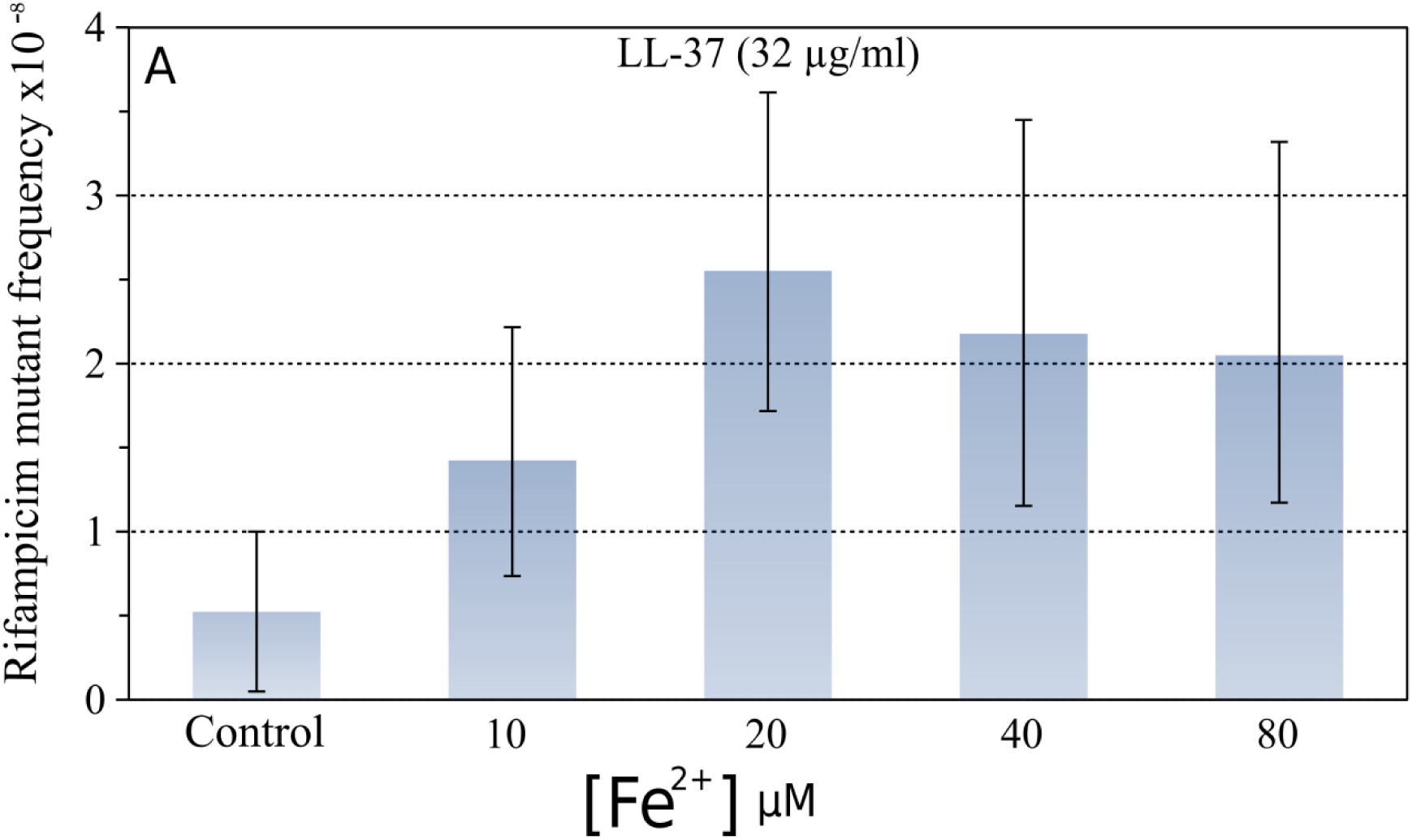
Different concentrations of Fe^2+^ increase mutant frequencies to rifampicin of *P. aeruginosa* PAO1 if LL-37 is present. The experiment was carried out at MIC50 for LL-37 (32 μg/ml). Error bars represent 95 % confidence intervals for mutant frequencies.

**Figure S4.**
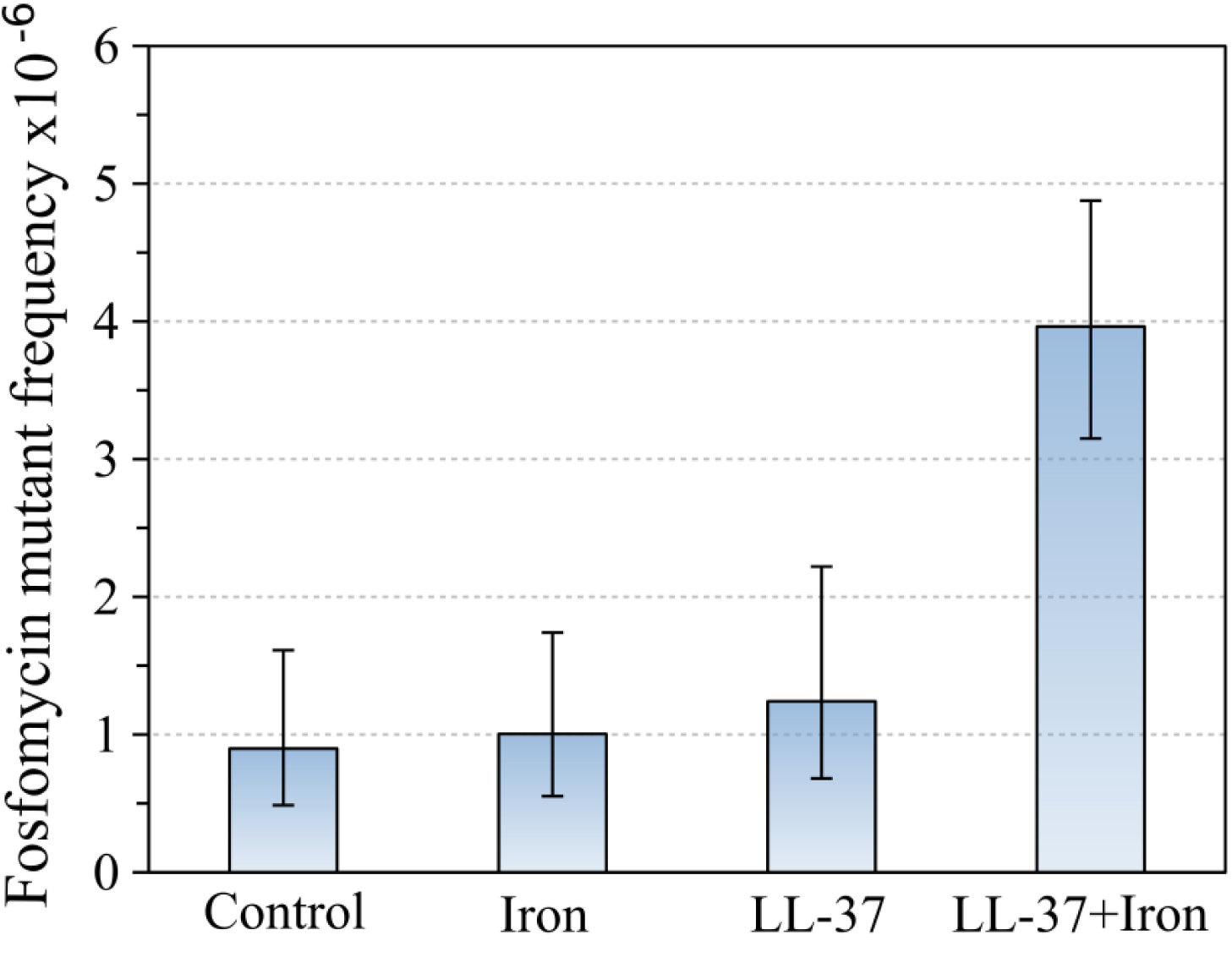
The combination of Fe^2+^ (40 μM) and LL-37 (32 μg/ml) increases the mutant frequency of *P. aeruginosa* PA14 to fosfomycin. Error bars represent 95 % confidence intervals for mutant frequencies.

**Figure S5.**
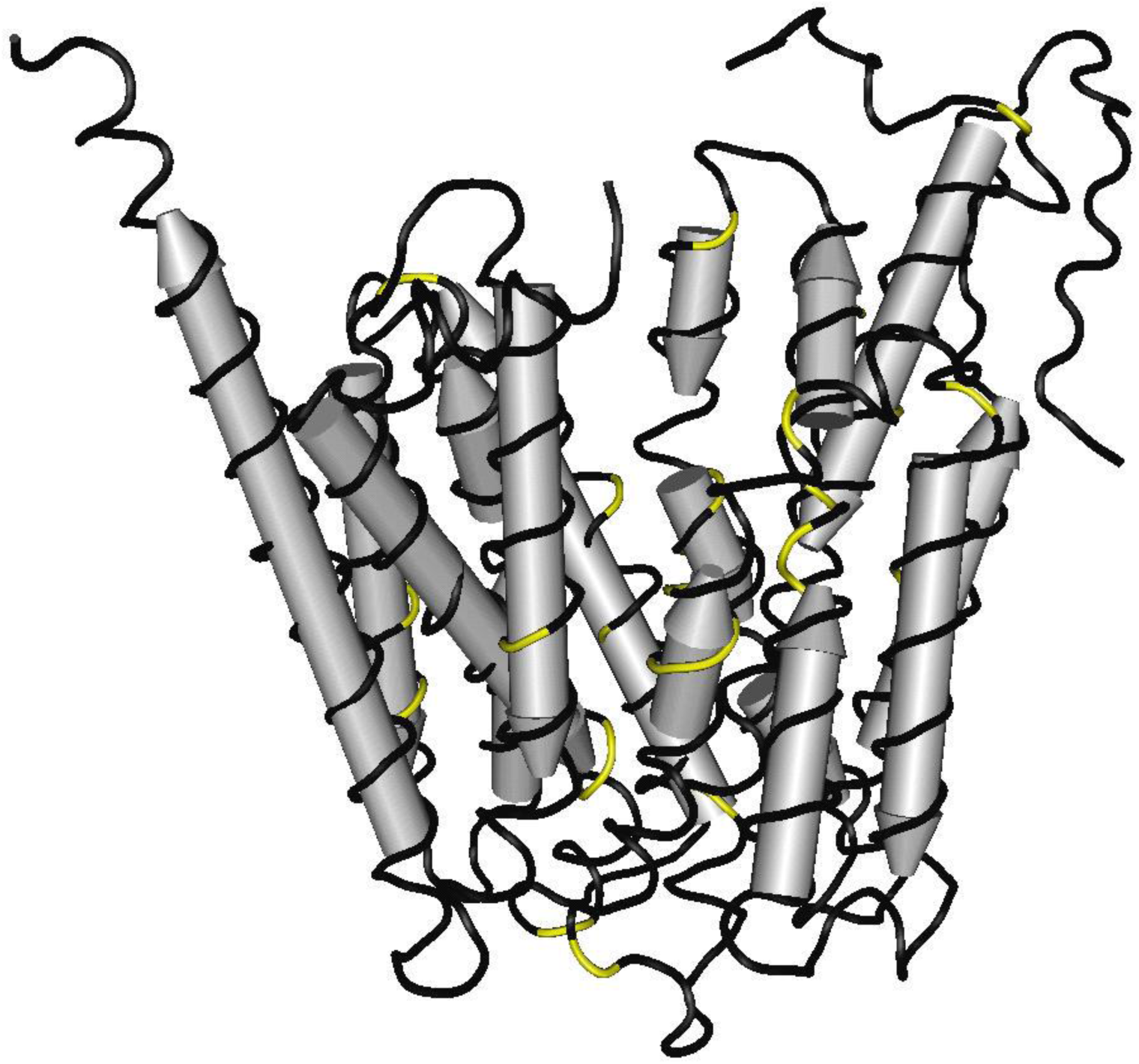
3D-structural model of GlpT. A homology model of GlpT was generated using Cn3D software by performing a BLASTP search using PA14 GlpT protein sequence as a query and mapping the resulting alignment against the experimentally determined *Escherichia coli* K-12 GlpT protein structure. Substitutions found in Fos-R mutants in all treatments are highlighted in yellow.

**Figure S6.** DNA alignment of the 1160 bp fragment of *glpT* from twenty Fos-R *P. aeruginosa* PA14 clones treated with iron, LL-37 and LL-37+Fe^2+^. The 52-1212 bp region relative to the A in the start ATG codon of the 1347bp-long *glpT* ORF.

**Figure S7.**
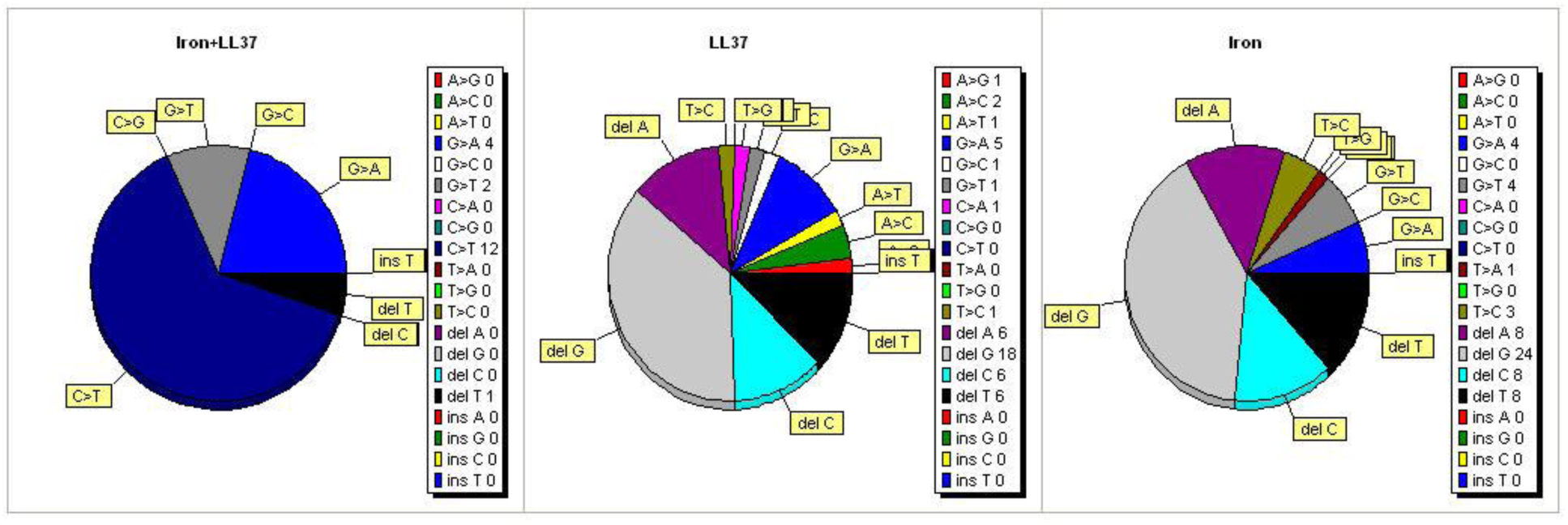
Mutation types in *glpT*. Prevalence of certain mutation types in iron, LL-37 and LL-37+Fe^2+^ treatments. C to T transitions was the most common type of nucleotide substitutions in LL-37+Fe^2+^ treatment, but not in two other treatments.

**Table S1.**
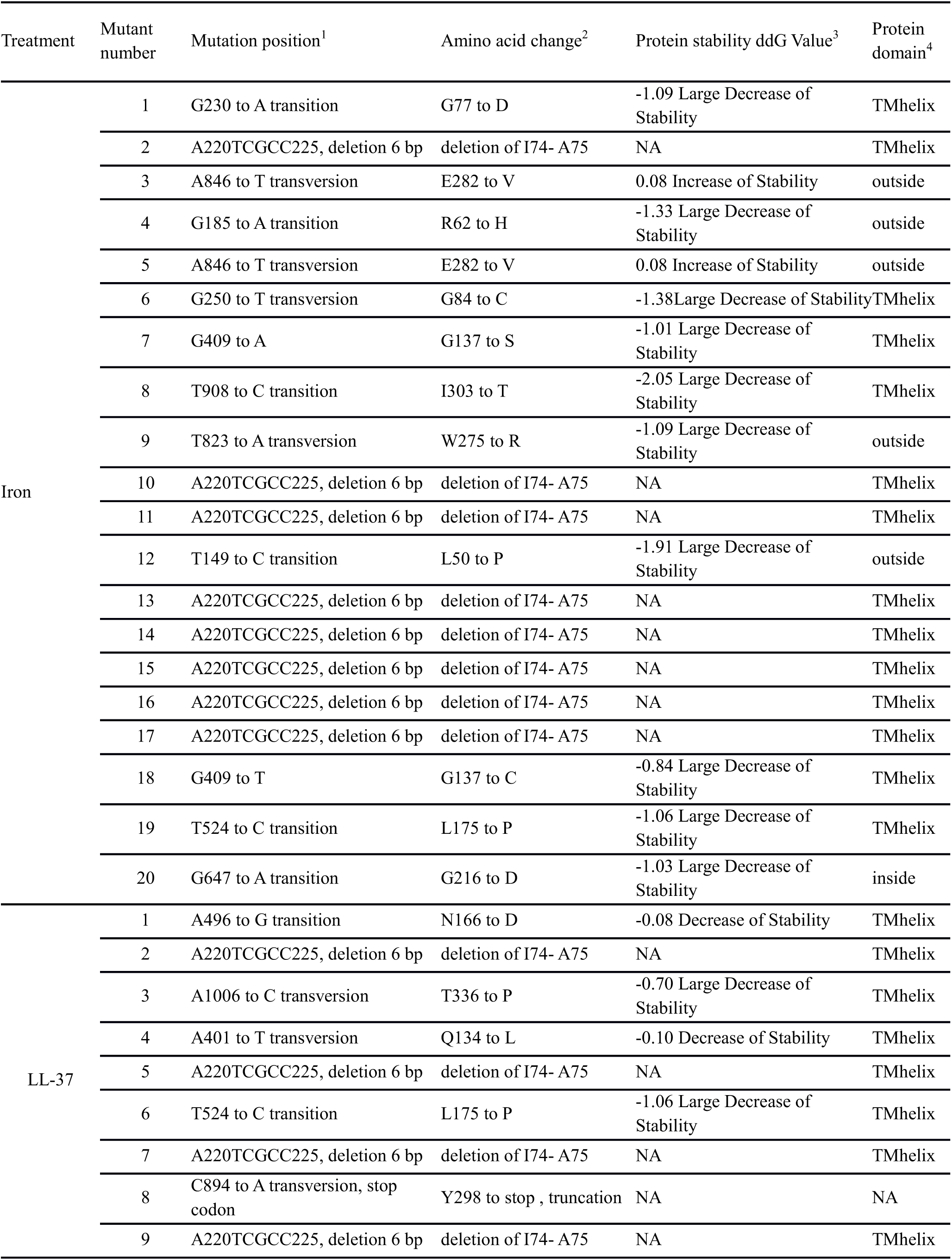
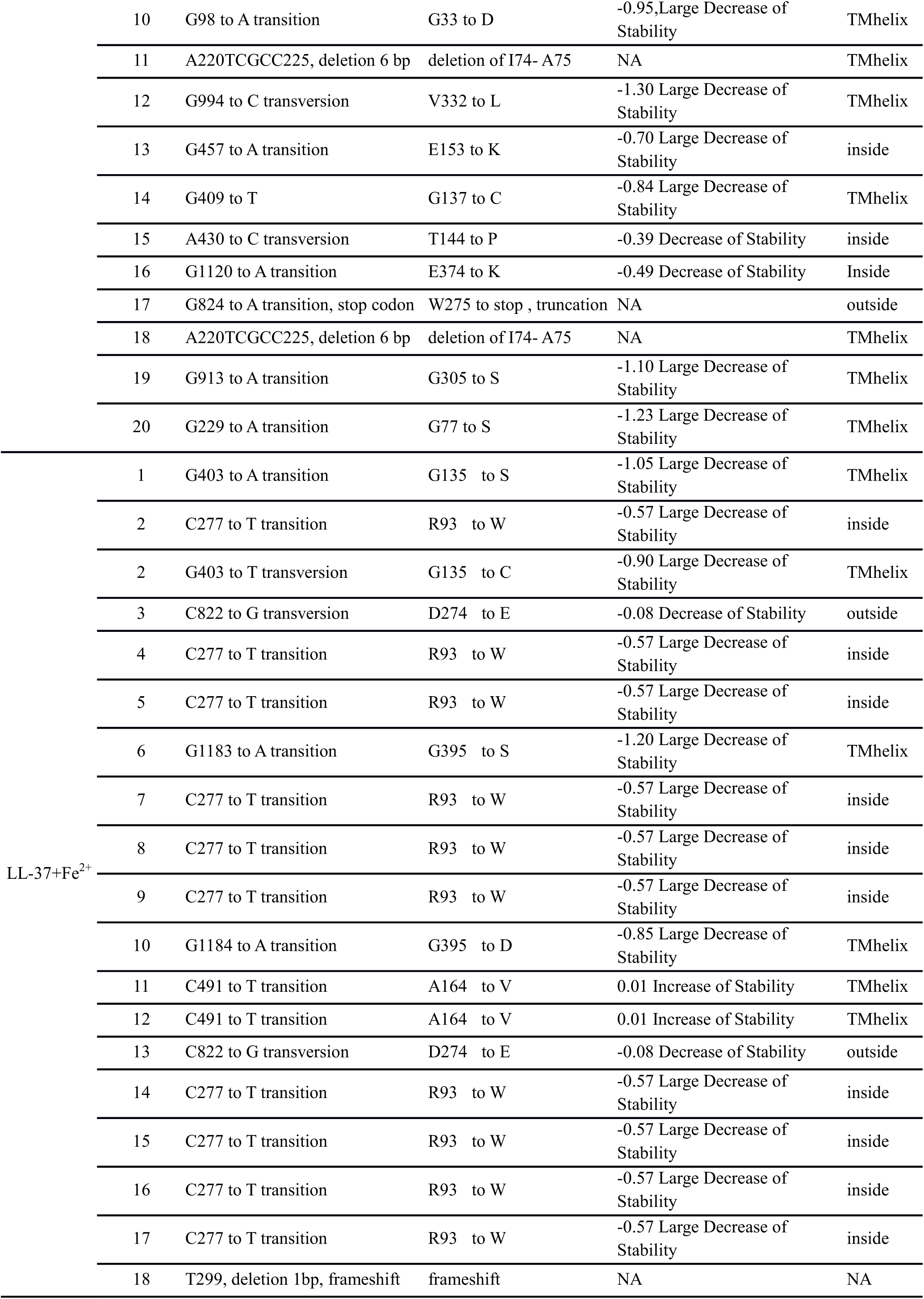
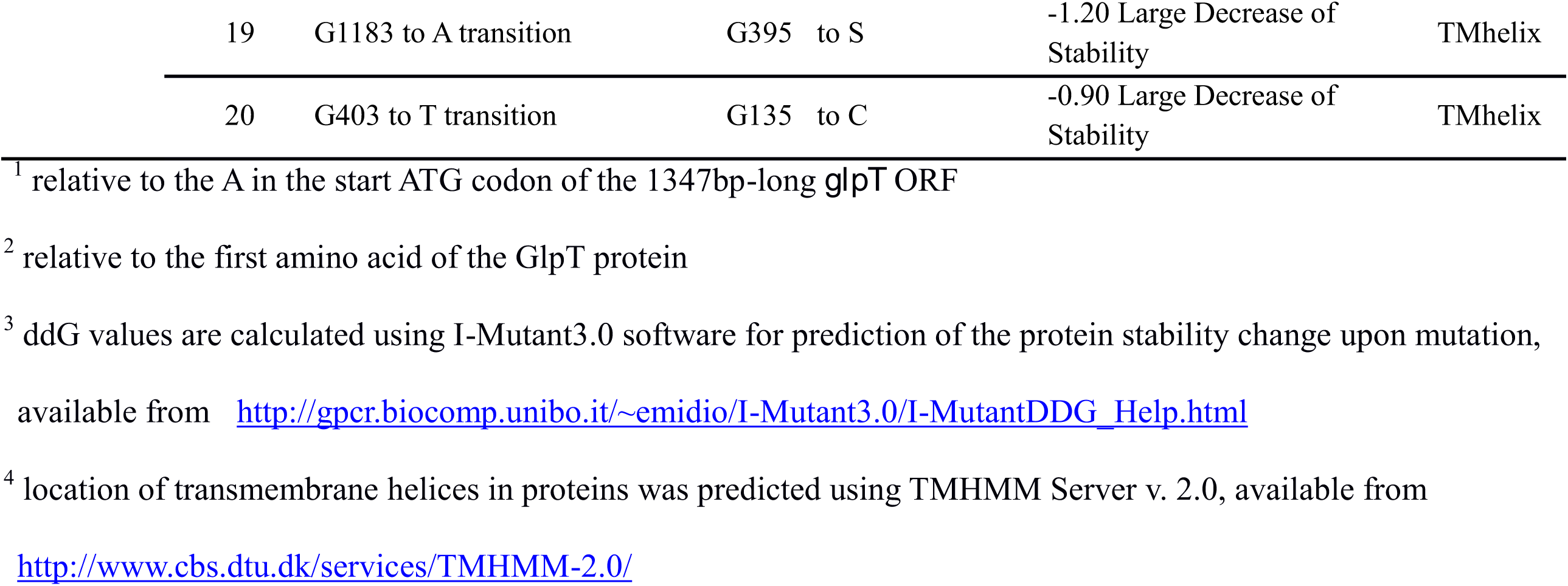
Mutations in *glpT* of Fos-R *P. aeruginosa* PA14 clones treated with iron, LL-37 and LL-37+Fe^2+^.

**Table S2.** Two-tailed Fisher’s exact-test of probability of having the same mutation in both LL-37 and iron treatments. H0 is rejected if *P*<0.05.

| Mutation site | Number of mutants per treatment |  | Two-tailed *P*-value | Interpretation |
| --- | --- | --- | --- | --- |
|  | LL-37 | Iron |  |  |
| G33 to D | 1 | 0 | > 0.9999 | The groups are not significantly different. |
| L50 to P | 0 | 1 | > 0.9999 | The groups are not significantly different. |
| R62 to H | 0 | 1 | > 0.9999 | The groups are not significantly different. |
| deletion of 6 bp I74 - A75 | 6 | 8 | 0.7411 | The groups are not significantly different. |
| G77 to D | 1 | 1 | > 0.9999 | The groups are not significantly different. |
| G84 to C | 0 | 1 | > 0.9999 | The groups are not significantly different. |
| R93 to W | 0 | 0 | > 0.9999 | The groups are not significantly different. |
| Q134 to L | 1 | 0 | > 0.9999 | The groups are not significantly different. |
| G135 to S | 0 | 0 | > 0.9999 | The groups are not significantly different. |
| G137 to S | 1 | 2 | > 0.9999 | The groups are not significantly different. |
| T299 frameshift | 0 | 0 | > 0.9999 | The groups are not significantly different. |
| T144 to P | 1 | 0 | > 0.9999 | The groups are not significantly different. |
| E153 to K | 1 | 0 | > 0.9999 | The groups are not significantly different. |
| N166 to D | 1 | 0 | > 0.9999 | The groups are not significantly different. |
| A164 to V | 0 | 0 | > 0.9999 | The groups are not significantly different. |
| L175 to P | 1 | 1 | > 0.9999 | The groups are not significantly different. |
| G216 to D | 0 | 1 | > 0.9999 | The groups are not significantly different. |
| D274 to E | 0 | 0 | > 0.9999 | The groups are not significantly different. |
| W275 to R | 1 | 1 | > 0.9999 | The groups are not significantly different. |
| E282 to V | 0 | 2 | 0.4872 | The groups are not significantly different. |
| Y298 to stop, truncation | 1 | 0 | > 0.9999 | The groups are not significantly different. |
| I303 to T | 0 | 1 | > 0.9999 | The groups are not significantly different. |
| G305 to S | 1 | 0 | > 0.9999 | The groups are not significantly different. |
| V332 to L | 1 | 0 | > 0.9999 | The groups are not significantly different. |
| T336 to P | 1 | 0 | > 0.9999 | The groups are not significantly different. |
| E374 to K | 1 | 0 | > 0.9999 | The groups are not significantly different. |
| G395 to S | 0 | 0 | > 0.9999 | The groups are not significantly different. |

**Table S3.** Two-tailed Fisher’s exact-test of probability of having the same mutation in both LL-37 and LL-37+Fe^2+^ treatments. H0 is rejected if *P*<0.05. Mutation hotspots that significantly different are highlighted.

| Mutation site | Number of mutants per treatment |  | Two-tailed *P*-value | Interpretation |
| --- | --- | --- | --- | --- |
|  | LL-37 | LL-37+Fe<sup>2+</sup> |  |  |
| G33 to D | 1 | 0 | > 0.9999 | The groups are not significantly different. |
| L50 to P | 0 | 0 | > 0.9999 | The groups are not significantly different. |
| R62 to H | 0 | 0 | > 0.9999 | The groups are not significantly different. |
| deletion of 6 bp I74 - A75 | 6 | 0 | 0.0202 | The groups are significantly different. |
| G77 to D | 1 | 0 | > 0.9999 | The groups are not significantly different. |
| G84 to C | 0 | 0 | > 0.9999 | The groups are not significantly different. |
| R93 to W | 0 | 10 | 0.0004 | The groups are significantly different. |
| Q134 to L | 1 | 0 | > 0.9999 | The groups are not significantly different. |
| G135 to S | 0 | 3 | 0.2308 | The groups are not significantly different. |
| G137 to S | 1 | 0 | > 0.9999 | The groups are not significantly different. |
| T299 frameshift | 0 | 1 | > 0.9999 | The groups are not significantly different. |
| T144 to P | 1 | 0 | > 0.9999 | The groups are not significantly different. |
| E153 to K | 1 | 0 | > 0.9999 | The groups are not significantly different. |
| N166 to D | 1 | 0 | > 0.9999 | The groups are not significantly different. |
| A164 to V | 0 | 2 | 0.4872 | The groups are not significantly different. |
| L175 to P | 1 | 0 | > 0.9999 | The groups are not significantly different. |
| G216 to D | 0 | 0 | > 0.9999 | The groups are not significantly different. |
| D274 to E | 0 | 2 | 0.4872 | The groups are not significantly different. |
| W275 to R | 1 | 0 | > 0.9999 | The groups are not significantly different. |
| E282 to V | 0 | 0 | > 0.9999 | The groups are not significantly different. |
| Y298 to stop, truncation | 1 | 0 | > 0.9999 | The groups are not significantly different. |
| I303 to T | 0 | 0 | > 0.9999 | The groups are not significantly different. |
| G305 to S | 1 | 0 | > 0.9999 | The groups are not significantly different. |
| V332 to L | 1 | 0 | > 0.9999 | The groups are not significantly different. |
| T336 to P | 1 | 0 | > 0.9999 | The groups are not significantly different. |
| E374 to K | 1 | 0 | > 0.9999 | The groups are not significantly different. |
| G395 to S | 0 | 3 | 0.2308 | The groups are not significantly different. |

**Table S4.** Two-tailed Fisher’s exact-test of probability of having the same mutation in both iron and LL-37+Fe^2+^ treatments. H0 is rejected if *P*<0.05. Mutation hotspots that significantly different are highlighted.

| Mutation site | Number of mutants per treatment |  | Two-tailed *P*-value | Interpretation |
| --- | --- | --- | --- | --- |
|  | Iron | LL-37+Fe<sup>2+</sup> |  |  |
| G33 to D | 0 | 0 | > 0.9999 | The groups are not significantly different. |
| L50 to P | 1 | 0 | > 0.9999 | The groups are not significantly different. |
| R62 to H | 1 | 0 | > 0.9999 | The groups are not significantly different. |
| deletion of 6 bp I74 - A75 | 8 | 0 | 0.0033 | The groups are significantly different. |
| G77 to D | 1 | 0 | > 0.9999 | The groups are not significantly different. |
| G84 to C | 1 | 0 | > 0.9999 | The groups are not significantly different. |
| R93 to W | 0 | 10 | 0.0004 | The groups are significantly different. |
| Q134 to L | 0 | 0 | > 0.9999 | The groups are not significantly different. |
| G135 to S | 0 | 3 | 0.2308 | The groups are not significantly different. |
| G137 to S | 2 | 0 | 0.4872 | The groups are not significantly different. |
| T299 frameshift | 0 | 1 | > 0.9999 | The groups are not significantly different. |
| T144 to P | 0 | 0 | > 0.9999 | The groups are not significantly different. |
| E153 to K | 0 | 0 | > 0.9999 | The groups are not significantly different. |
| N166 to D | 0 | 0 | > 0.9999 | The groups are not significantly different. |
| A164 to V | 0 | 2 | 0.4872 | The groups are not significantly different. |
| L175 to P | 1 | 0 | > 0.9999 | The groups are not significantly different. |
| G216 to D | 1 | 0 | > 0.9999 | The groups are not significantly different. |
| D274 to E | 0 | 2 | 0.4872 | The groups are not significantly different. |
| W275 to R | 1 | 0 | > 0.9999 | The groups are not significantly different. |
| E282 to V | 2 | 0 | > 0.9999 | The groups are not significantly different. |
| Y298 to stop, truncation | 0 | 0 | > 0.9999 | The groups are not significantly different. |
| I303 to T | 1 | 0 | > 0.9999 | The groups are not significantly different. |
| G305 to S | 0 | 0 | > 0.9999 | The groups are not significantly different. |
| V332 to L | 0 | 0 | > 0.9999 | The groups are not significantly different. |
| T336 to P | 0 | 0 | > 0.9999 | The groups are not significantly different. |
| E374 to K | 0 | 0 | > 0.9999 | The groups are not significantly different. |
| G395 to S | 0 | 3 | 0.2308 | The groups are not significantly different. |

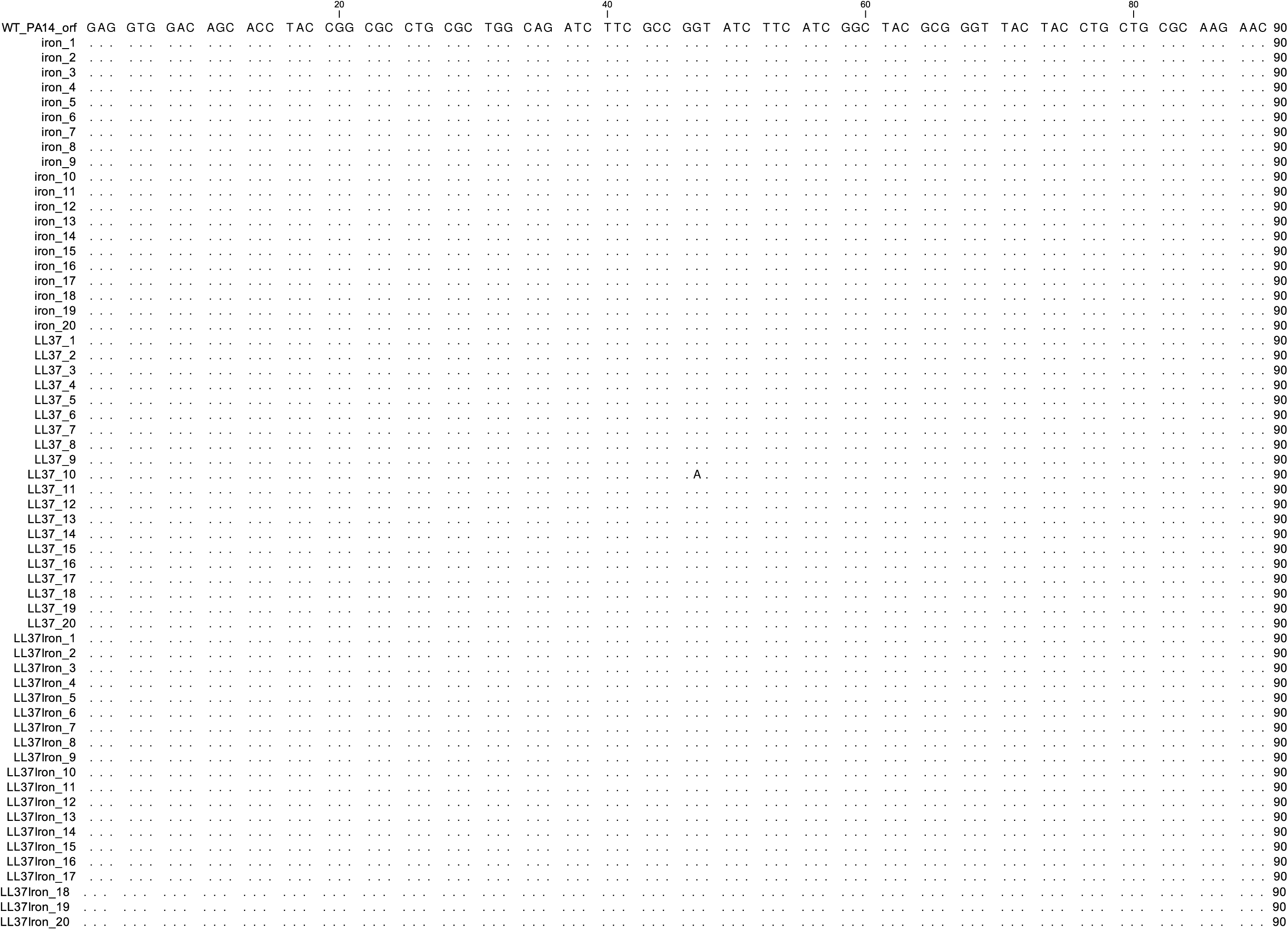

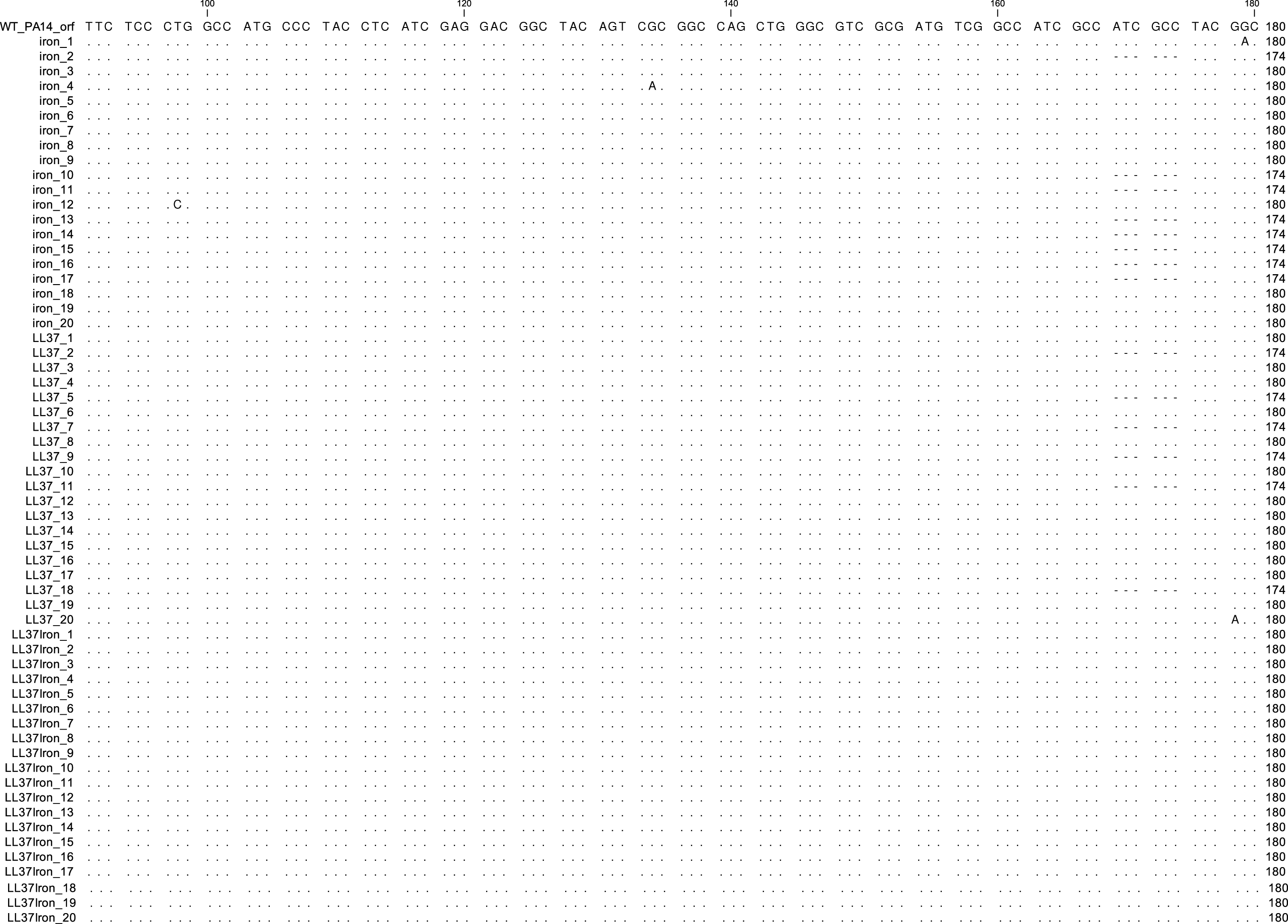

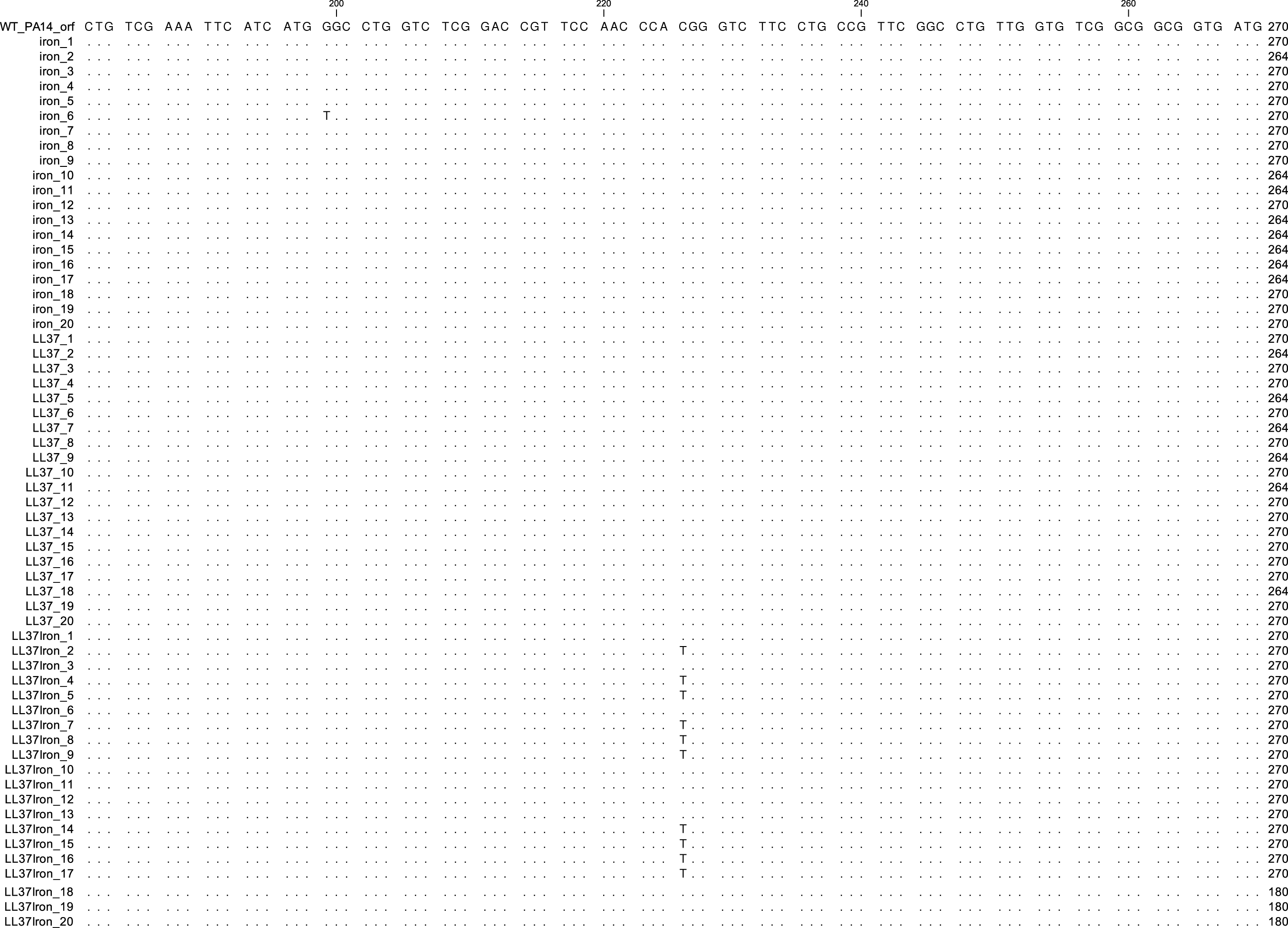

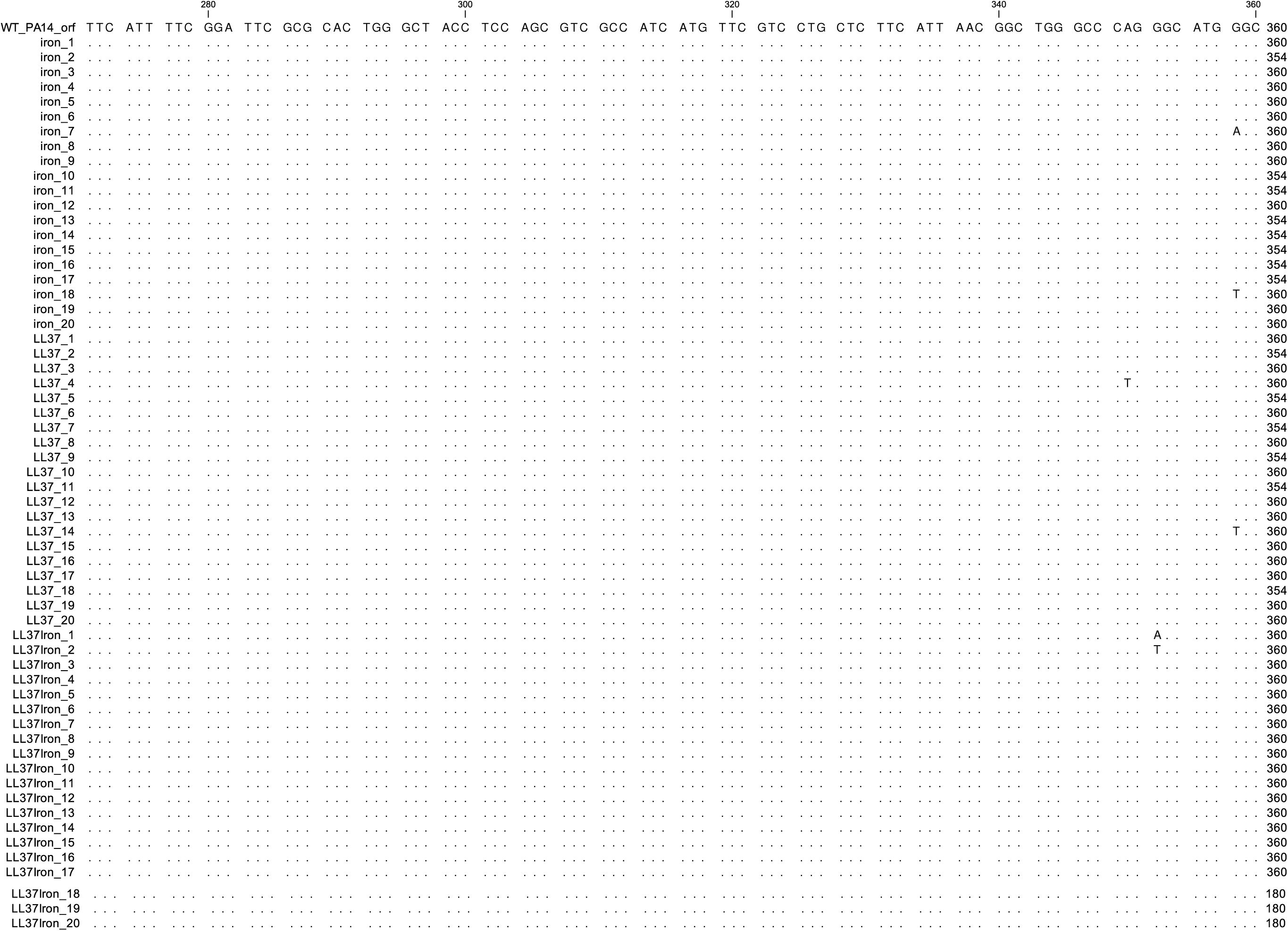

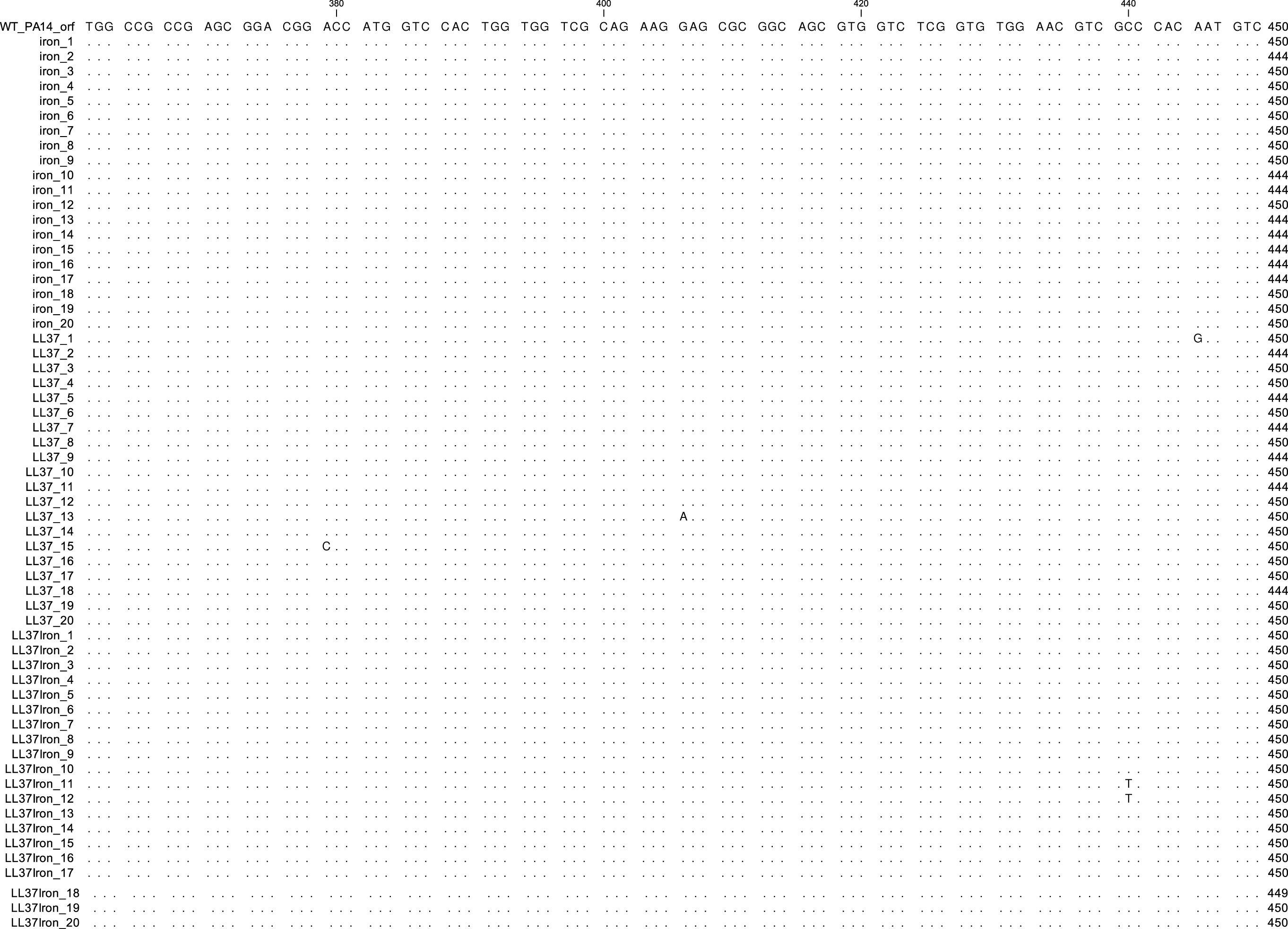

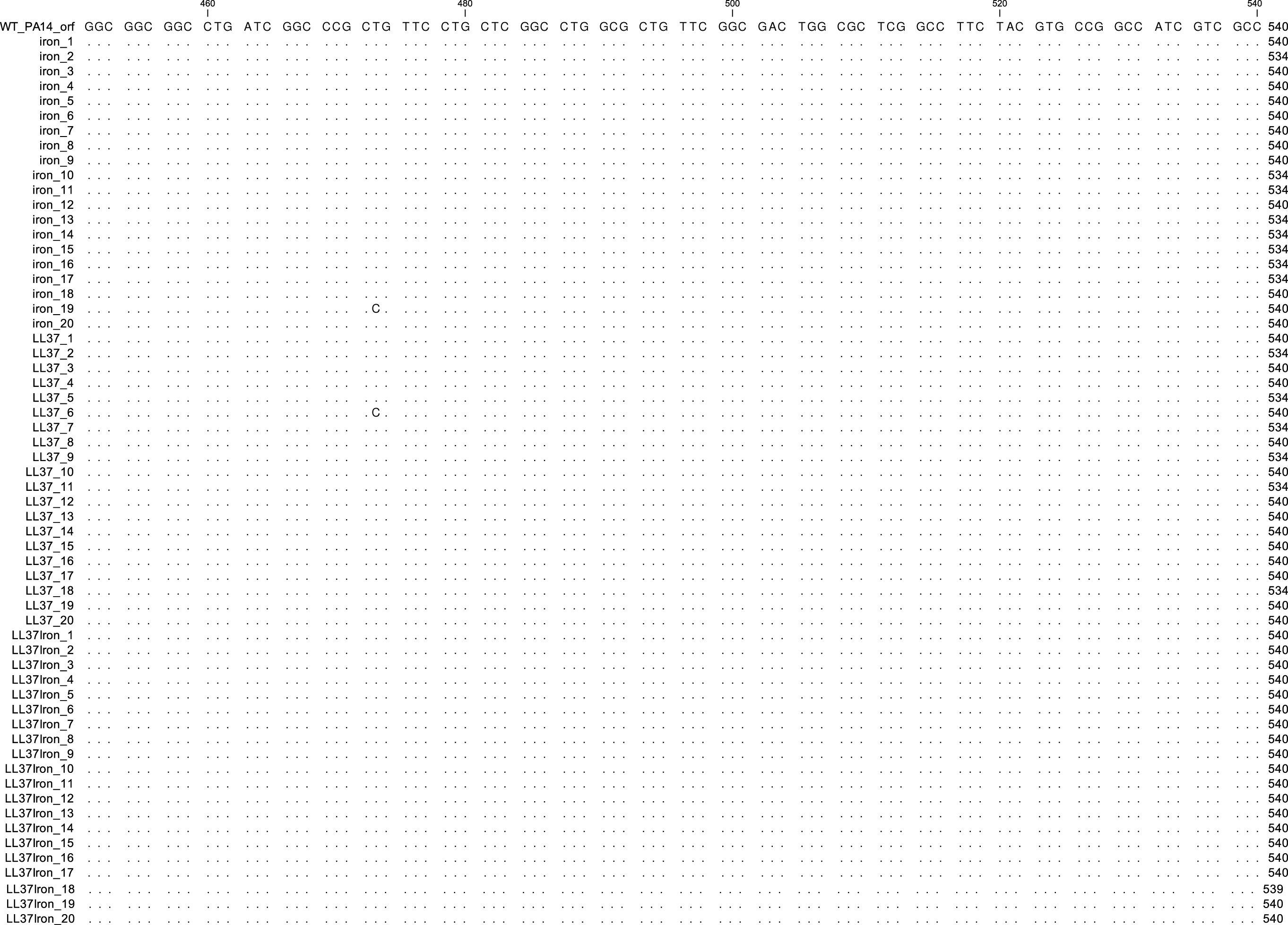

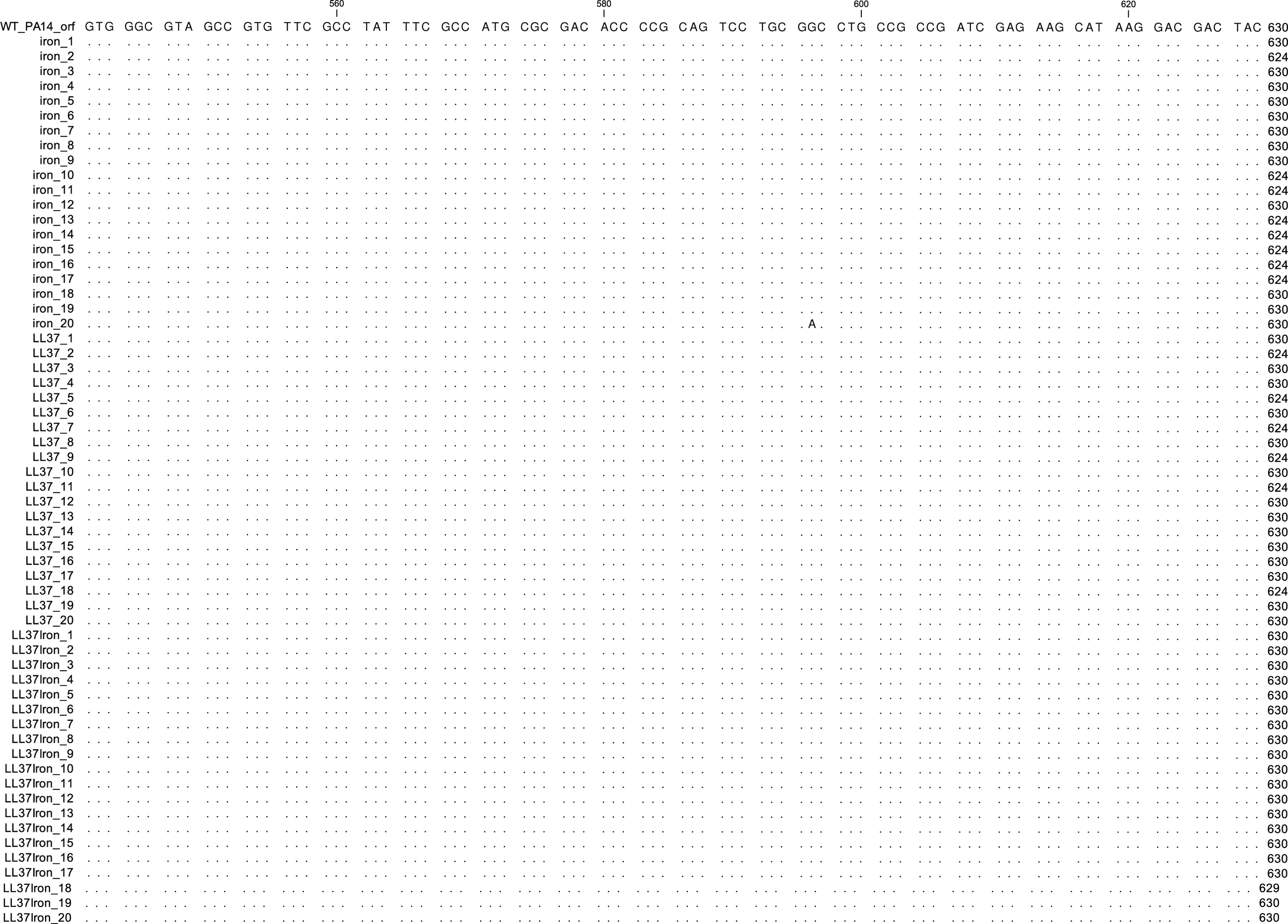

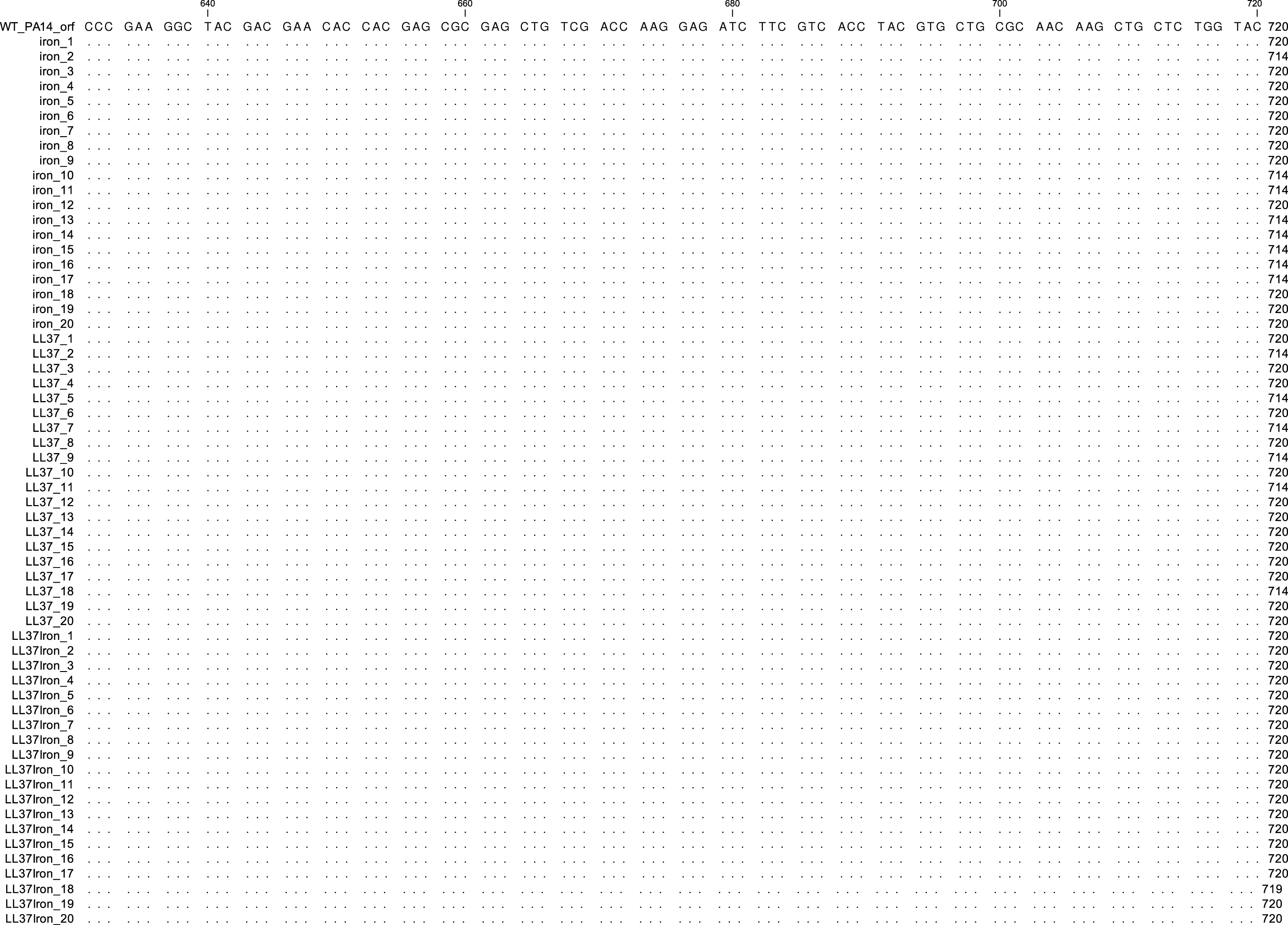

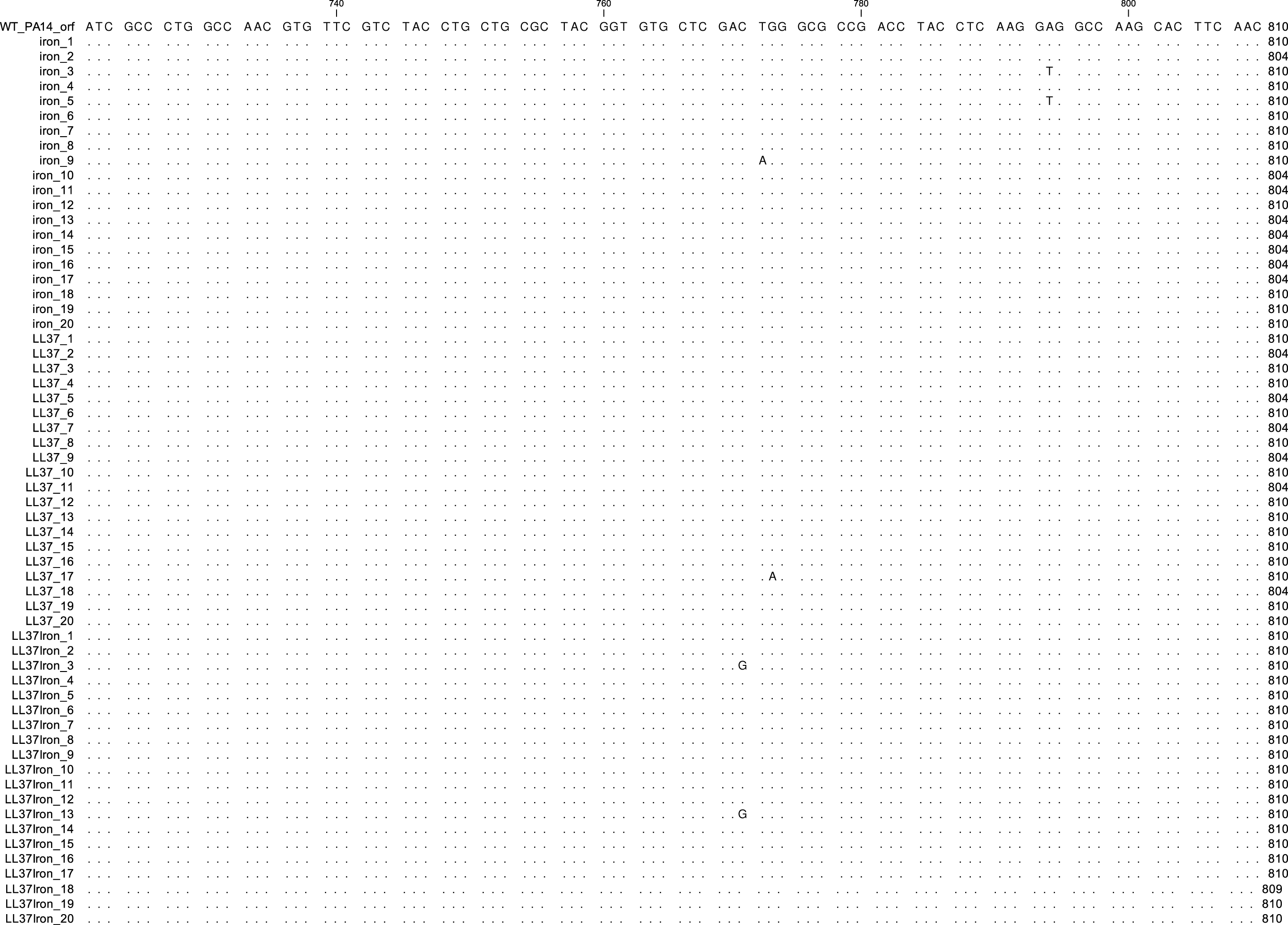

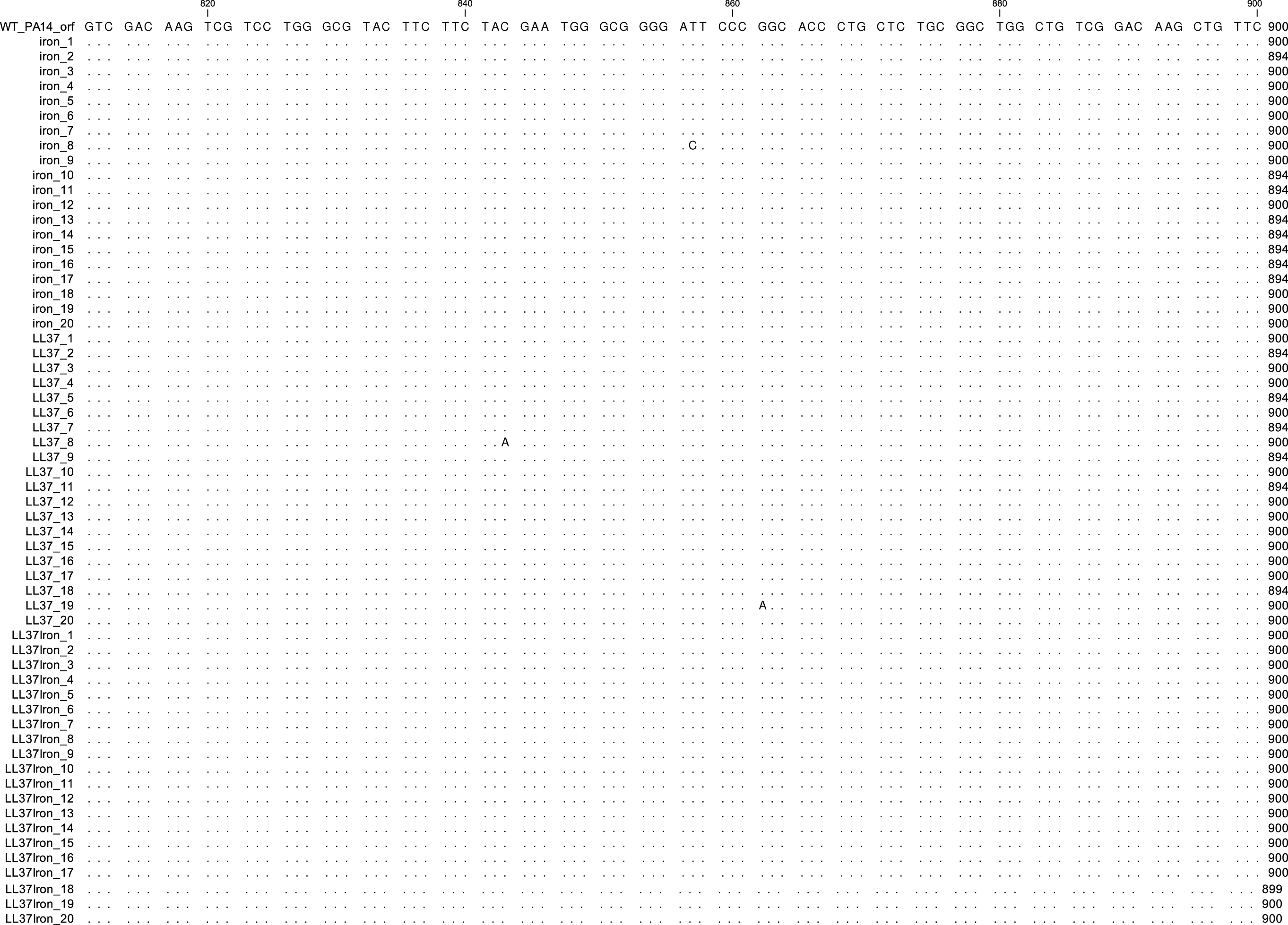

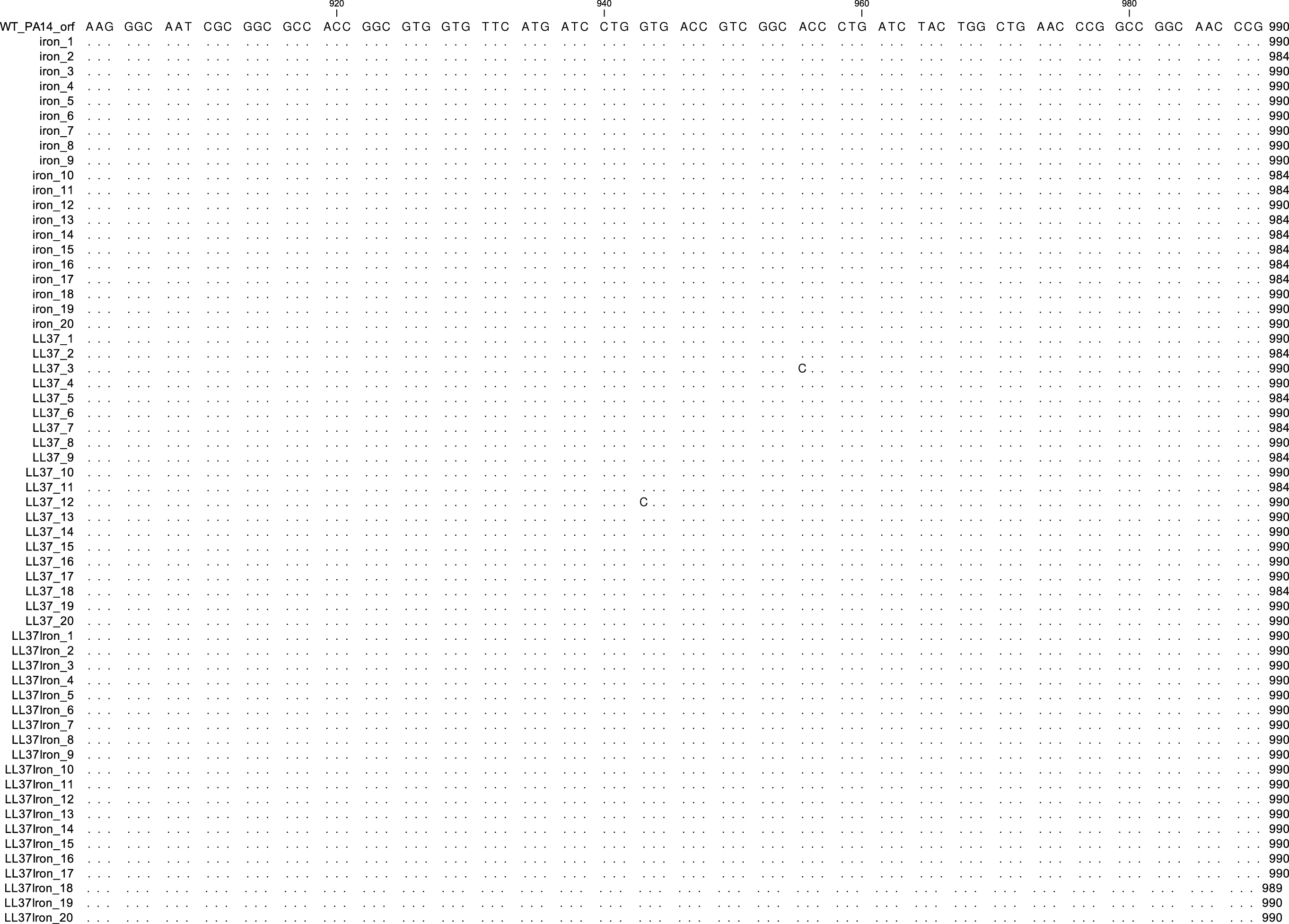

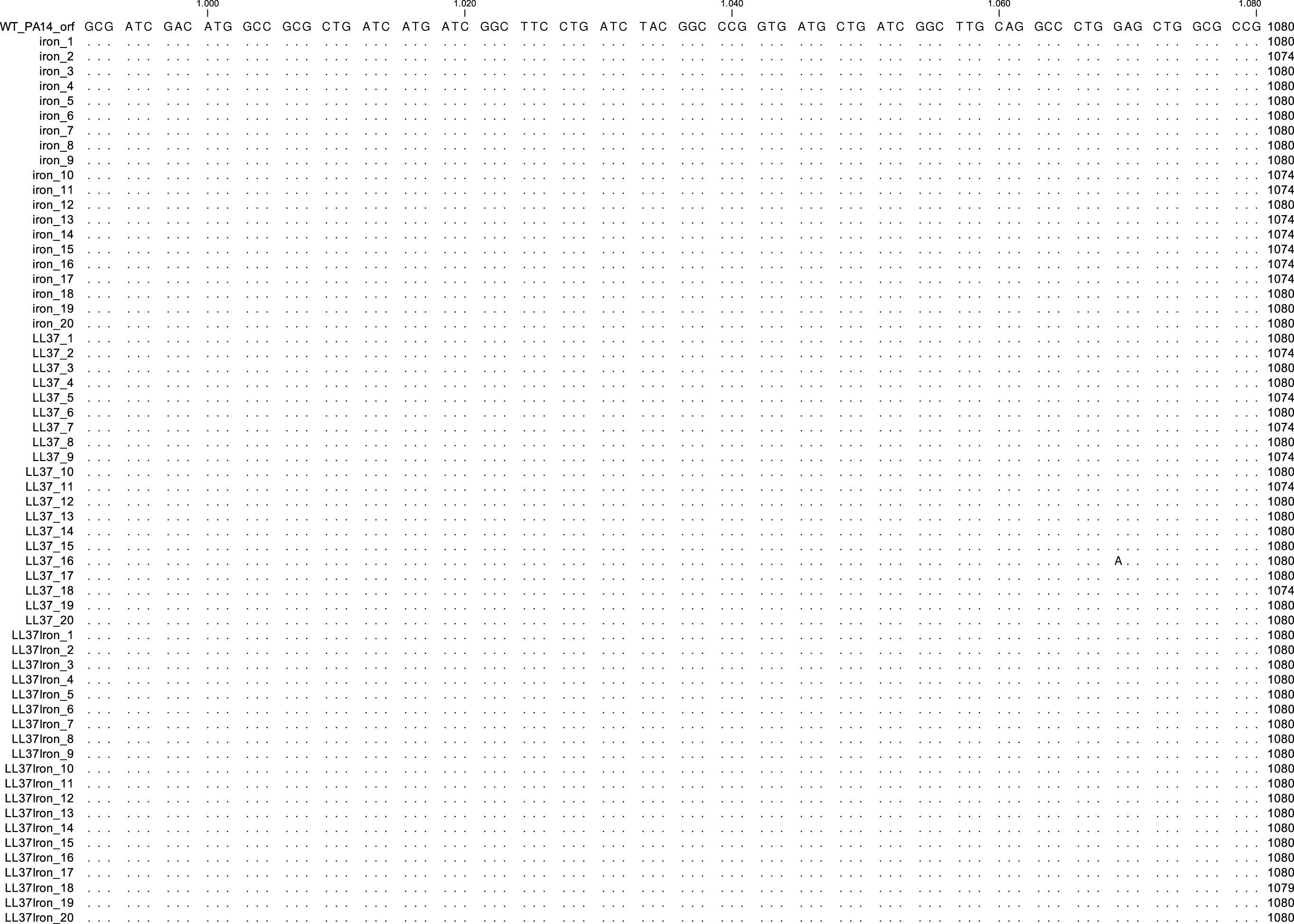

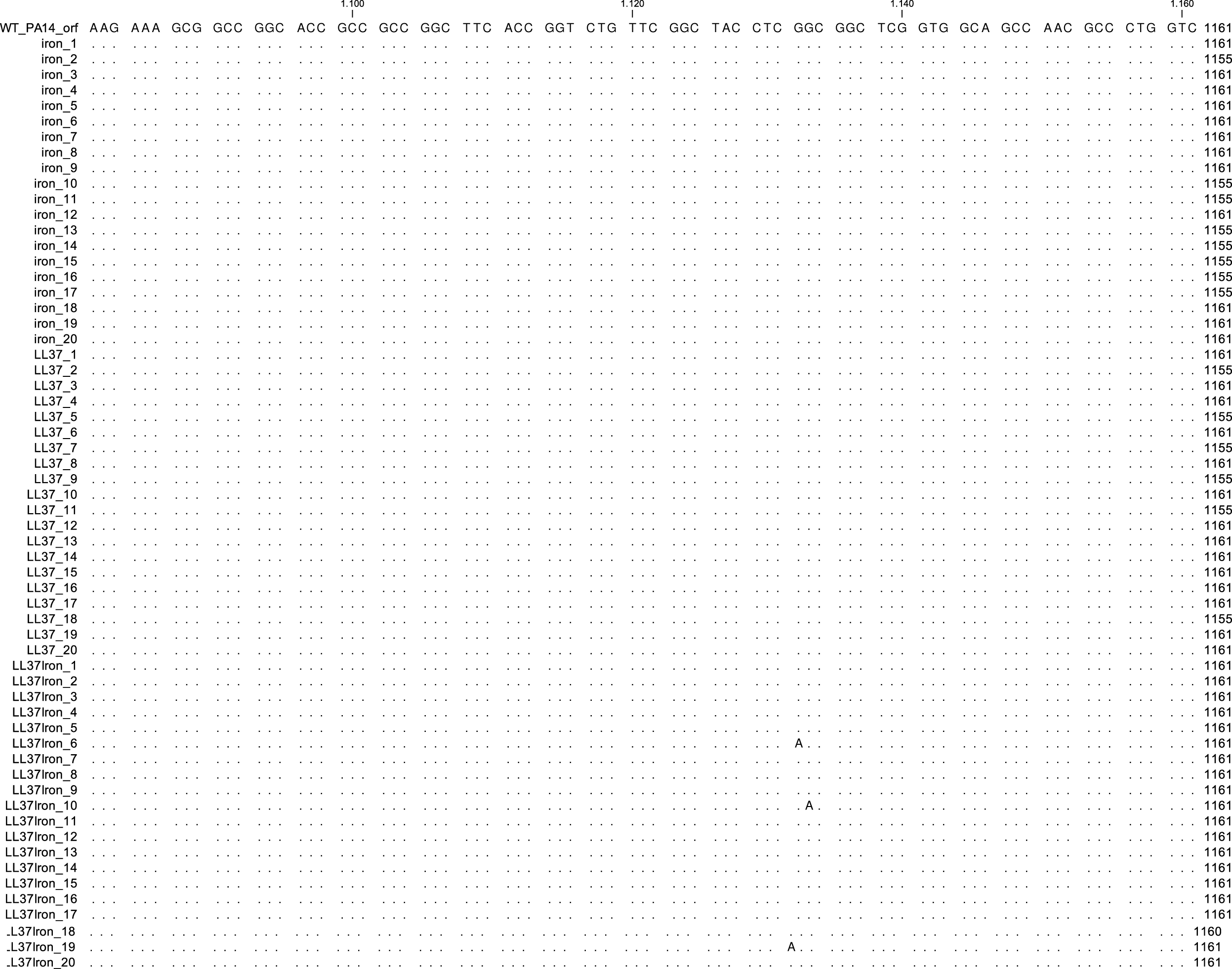

